# How does the mode of evolutionary divergence affect reproductive isolation?

**DOI:** 10.1101/2022.03.08.483443

**Authors:** Bianca De Sanctis, Hilde Schneemann, John J. Welch

## Abstract

When divergent populations interbreed, the outcome will be affected by the genomic and phenotypic differences that they have accumulated. In this way, the mode of evolutionary divergence between populations may have predictable consequences for the fitness of their hybrids, and so for the progress of speciation. To investigate these connections, we present a new analysis of hybridization under Fisher’s geometric model, making few assumptions about the allelic effects that differentiate the hybridizing populations. Results show that the strength and form of postzygotic reproductive isolation (RI) depend on just two properties of the evolutionary changes, which we call the “total amount” and “net effect” of change, and whose difference quantifies the similarity of the changes at different loci, or their tendency to act in the same phenotypic direction. It follows from our results that identical patterns of RI can arise in different ways, since different evolutionary histories can lead to the same total amount and net effect of change. Nevertheless, we show how these estimable quantities do contain some information about the history of divergence, and that – thanks to Haldane’s Sieve – the dominance and additive effects contain complementary information.

**Impact Summary:** When populations of animals or plants evolve differences in their genomes or traits, the nature of the differences will help to determine whether they can continue to interbreed. For example, the hybrid offspring may be infertile, or unlikely to survive to reproductive age, meaning that the two populations remain distinct from one another even after mating. However, in some cases the hybrids may be more fertile than their parents or have some other reproductive advantage. In this study, we use a mathematical model to relate hybrid fitness to the evolved differences separating the parents. We find that the outcome depends on just two properties of these differences, which capture the “total amount” and the “net effect” of evolutionary change. We then show that different evolutionary divergence scenarios or modes can lead to the exact same hybrid fitness. On the other hand, we can still make some inferences about the history of divergence by observing certain properties of hybrid fitness. Determining the relationship between hybrid fitness and the mode of evolutionary divergence will help to understand how new species form, to plan conservation interventions such as moving individuals between isolated populations to increase their adaptive potential, and to understand how existing species might interact when their habitats overlap, for example due to climate change or other human impacts.

## Introduction

Genomic and phenotypic differentiation between populations are a major cause of reproductive isolation (RI), preventing hybrids from forming, or reducing their fitness when they do form. However, differentiation can also be a source of adaptive variation, if hybrids contain new fit combinations of traits or alleles, or act as conduits passing existing combinations from one population to another (Arnold and Hodges, 1995; Edmands, 1999, 2002; Coyne and Orr, 2004; Bierne et al., 2013; Schluter and Conte, 2009; Bernardes et al., 2017; Coughlan and Matute, 2020).

Which of these outcomes actually takes place must depend on the types of phenotypic and genomic differences that have accumulated before the hybrids form. A fundamental challenge in evolutionary biology is to understand the connections between the mode of evolutionary divergence, the type of differences that accrue, and the outcomes of subsequent hybridization. This can be framed in two ways: what can we learn about the (unobserved) history of parental divergence by observing their hybrids? (Lande, 1981; Welch, 2004; Schneemann et al., 2020; Fraser, 2020); and conversely, which divergence scenarios will predictably lead to RI? (Coyne and Orr, 2004). What, for example, are the respective roles of large-versus small-effect mutations in causing RI, and what are the roles of natural selection versus genetic drift (Lynch, 1991; Coyne and Orr, 2004; Jezkova et al., 2013; Satokangas et al., 2020; Moran et al., 2021; Clo et al., 2021)? All of these questions are essential for understanding the opposing processes of speciation and adaptive introgression (Abbott et al., 2013), and predicting the outcomes of novel hybridizations, including those that are human-mediated (Genovart, 2008; Chan et al., 2019).

One tool to address these questions is Fisher’s geometric model. This is a mathematical model of selection acting on quantitative traits (Fisher, 1930, Ch. 2), and has been used to understand both phenotypic data, e.g., QTL for traits involved in adaptive divergence (Orr, 1998), and fitness data. In the latter case, the phenotypic model need not be treated literally, but is a simple way of generating a fitness landscape (Martin and Lenormand, 2006; Martin, 2014). Both uses of the model have been applied to hybrids (Lande, 1981; Mani and Clarke, 1990; Barton, 2001; Chevin et al., 2014; Fraïsse et al., 2016; Simon et al., 2018; Yamaguchi and Otto, 2020; Schneemann et al., 2020; Thompson et al., 2021; Schneemann et al., 2022).

Most importantly here, the model allows us to consider the effects in hybrids of evolutionary changes of different sizes, and which were driven by different evolutionary processes (Hartl and Taubes, 1996; Orr, 1998; Chevin et al., 2014; Simon et al., 2018; Schneemann et al., 2020). However, previous analytical results for diploids (Schneemann et al., 2020) depended on strong assumptions about the genetic differentiation, such as no variation within the parental lines, normality and universal pleiotropy among the fixed effects, and statistical independence among traits. Furthermore, the earlier results describe the overall strength of RI in terms of a single fitted parameter, whose relationship to the process of evolutionary divergence remained obscure.

In this paper, we extend previous work on Fisher’s geometric model in two ways. First, by combining and generalizing previous work by several authors (Lande, 1981; Chevin et al., 2014; Simon et al., 2018; Schneemann et al., 2020, 2022), we give results for the expected fitness of hybrids between diploid populations, applying to all classes of hybrid, and allowing for variation within the hybridizing populations, and alleles with arbitrary additive and dominance effects. Second, we show how some key quantities that appear in the results relate transparently to the history of divergence between the parental populations.

## Results

### The phenotypic model and fitness landscape

Under Fisher’s geometric model, the fitness of any individual depends solely on its values of *n* quantitative traits. The trait values for an individual can be collected in an *n*-dimensional vector **z** = (*z*_i_,…, *z_n_*); and its fitness, *w*, depends on the Euclidean distance of this phenotype from an optimum **o** = (*o*_1_,…, *o_n_*), whose value is determined by the current environment. We will assume the simplest form of the model, where the log fitness declines with the square of the distance:

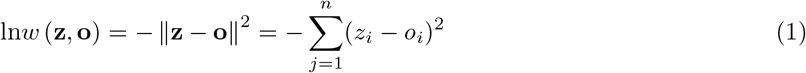

This model can be derived either exactly, or approximately, from a wide class of more complicated fitness functions (Martin, 2014; Schneemann et al., 2020), and in these latter cases, only a few, if any of the *n* traits, need to be identified with real quantitative traits that might be measured in the field. Results can also be applied if fitness declines more rapidly with distance from the optimum. For example, if ln*w* = – ║**z** – **o**║^*k*^ (Fraïsse et al., 2016; Simon et al., 2018; Fraïsse and Welch, 2019) then results below could be applied directly to the scaled log fitness (–ln*w*)^2/*k*^ = ║**z** – **o**║^2^.

### Characterizing parental divergence, and describing hybrids

We will consider hybrids between two diploid parental populations, denoted P1 and P2. We will assume that individuals in these populations vary at *D* biallelic loci, and that the allele frequencies might vary between populations, which includes the case when an allele is fixed in one population and absent in the other. If we (arbitrarily) choose one allele at each locus to be the focal allele, then the frequency of the focal allele at locus *i* = 1,…, *D* is denoted as *q*_P1,*i*_ (*q*_P2,*i*_) in population P1 (P2). We now make the key simplifying assumptions that (1) there are no statistical associations between alleles within the parental populations, so that both P1 and P2 are at Hardy-Weinberg and linkage equilibrium at all D loci, and (2) there is no phenotypic epistasis between the allelic effects.

With these assumptions, the differences in the trait means between P1 and P2 can be written as the sum of contributions from each of the *D* loci. As such, for any trait *j* = 1,…, *n*, the difference in trait means can be written

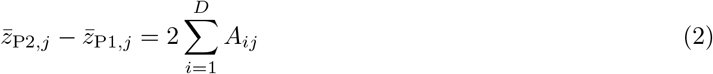

where the factor 2 follows from diploidy. A simple consequence of eq. 2 is that the phenotypic differentiation between the parental populations can be described as a chain of effects in *n*-dimensional phenotypic space. Figure 1A shows an illustrative example with *n* = 2 traits, affected by changes at *D* = 5 loci. Here, the black arrows represent the 2*A_ij_*, connecting the trait means of P1 and P2, or the centroids of the clouds of points that would represent the two parental populations. Each 2*A_ij_* describes the diploid effect on trait *j* of changing the allele frequency at locus *i* from *q*_P1,*i*_ to *q*_P2,*i*_.

**Figure 1:**
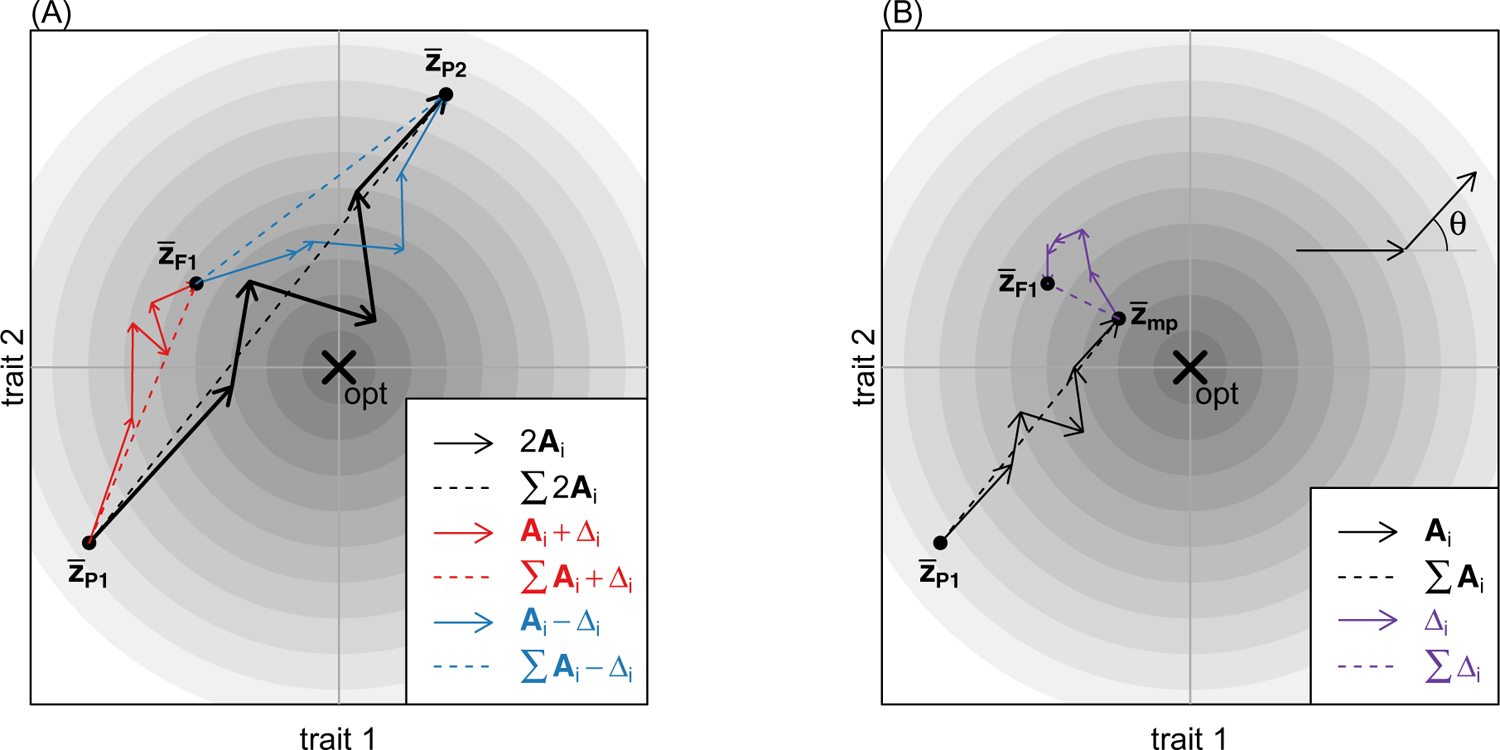
The key quantities that determine hybrid mean log fitness under Fisher’s geometric model. The fitness of any given phenotype is determined by its distance from some optimum phenotype, as determined by the current environment. This optimum and fitness landscape is illustrated, for *n* = 2 traits, by the cross and contour lines. **(A)**: The diploid parental populations, P1 and P2, are each characterized by mean phenotypic values, 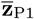 and 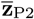, and the difference between these points are due to allele frequencies changes at *D* = 5 loci, each affecting one or more of the traits. The diploid changes associated with each locus are represented by the black arrows, whose components are denoted 2*A_ij_* for the diploid change to the *j*^th^ trait due to the *i*^th^ locus. The model allows for phenotypic dominance, so that the differences between the trait means of the parents, and the initial F1 cross, also involve dominance effects, denoted as Δ_*ij*_ for the change to the *j*^th^ trait due to the *i*^th^ locus. **(B)**: the additive (black) and dominance (purple) effects can also decomposed into chains of differences linking the P1 or F1 trait means to the mid-parental trait mean 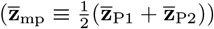. The mean log fitness of an arbitrary hybrid is affected by the *total amount of evolutionary change* (the sum of squared lengths of the arrows in a chain), and by the *net effect of the evolutionary change* (the squared lengths of the dotted lines). See text for full details.

We can also relate the *A_ij_* to the parental allele frequencies and the size of the phenotypic effect, as represented by the Fisherian average effect of a substitution (e.g. Lynch and Walsh, 1998, Ch. 4). In particular, we show in the Methods that

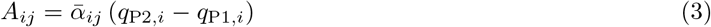

where 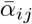 is the average effect of a substitution at locus *i* on trait *j* (e.g. Lynch and Walsh, 1998, eq. 4.10b), averaged across the two parental populations.

When there is phenotypic dominance (Lynch and Walsh, 1998, Ch. 4, Schneemann et al., 2022) we also need to account for the dominance deviations associated with allele frequency changes. We can do this by considering the mean phenotype in the initial F1 cross between P1 and P2, in which all loci in all individuals carry one P1-derived allele and one P2-derived allele. We show in the Methods that the difference in trait means between the F1, and the two parental populations can be written as

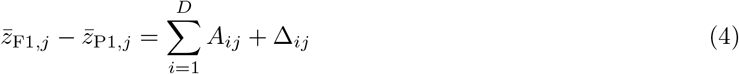

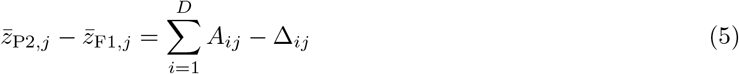

where

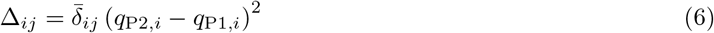

and 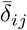 is the dominance deviation of a substitution at locus *i* on trait *j* averaged across the two parental populations. The differences between the parental and F1 trait means can also be represented as chains of effects, and this is illustrated by the red and blue arrows in Figure 1A. Moreover, we can separate out the additive and dominance effects by considering the differences between the F1 and the midparental mean phenotypes, defined as 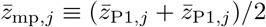.

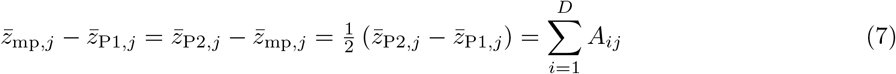

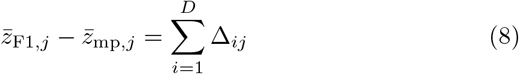

The two resulting chains are illustrated in Figure 1B.

The arguments above for the F1 cross generalize to an arbitrary hybrid (say, an F2 or a backcross). Hybrid genomes can be characterized in a number of different ways. In the main text, we will consider results for crosses, assuming free recombination among the *D* loci, and that no linkage disequilibrium has accumulated due to selection on early generation hybrids (see Lynch and Walsh, 1998 Ch. 9, and Schneemann et al., 2020 for some generalizations). In this case, hybrid genomes can be described solely in terms of their hybrid index, *h* (defined as the probability that a randomly chosen allele in the hybrid derives from parental line P2), and their inter-class heterozygosity, *p*_12_ (defined as the probability that a randomly chosen locus carries one allele of P1 origin and one allele of P2 origin). Results in the main text treat *h* and *p*_12_ as probabilities determined by the crossing scheme, and which apply to all loci independent of their allelic effects. In Appendix 1 we report equivalent results for sequenced genomes with known patterns of ancestry, such that *h* and *p*_12_ are known proportions. In either case, our aim is to calculate the expected fitness of a hybrid, conditional on *h* and *p*_12_. When we take expectations, they will be over the particular loci that are in any given ancestry state. We then determine how this result depends on properties of the additive and dominance effects. These will be collected in *D* × *n* - dimensional matrices, denoted **A** = (*A_ij_*) and **Δ** = (Δ*_ij_*), and treated as fixed observations, rather than random variables.

### Expected log fitness of a hybrid

Given the model described above, the expected log fitness of an arbitrary cross can be determined from the expected means and variances of its *n* traits.

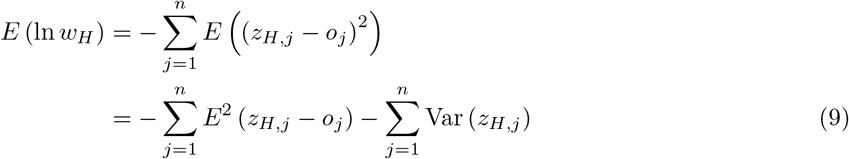

In the Methods, we show that each of the two terms in eq. 9 can be written as the sum of six terms, weighted by the same six combinations of *h* and *p*_12_. All 12 of these terms are shown in Table 1, where we introduce the notation

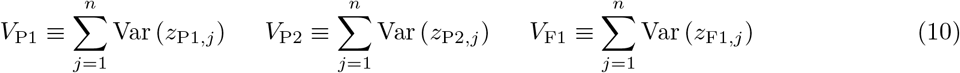

to denote the sum of the phenotypic variances over the *n* traits in a given population. We also introduce two new functions of *D* × *n* - dimensional matrices

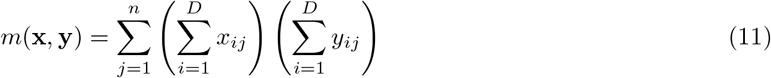

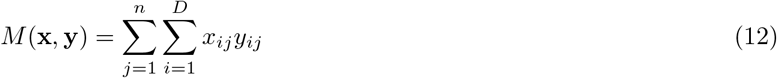

whose meanings we discuss below. The expected log fitness of any hybrid with a given value of *h* and *p*_12_ (eq. 9) is equal to the sum of the twelve terms in the second and third columns of Table 1, as weighted by their coefficients in the first column. Examining these terms, it follows that the expected log fitness depends on both properties of the parental populations (see top two rows of Table 1), and properties of the initial F1 cross (see third row of Table 1), plus properties of the additive and dominance effects, as captured by the functions *m*(·, ·) and *M*(·, ·) (see the bottom three rows of Table 1).

**Table 1:** Components of expected log hybrid fitness

| Coefficient | $-\sum_{j=1}^n E^2(z_H - o_j)$ | $-\sum_{j=1}^n \text{Var}(z_{H,j})$ |
| --- | --- | --- |
| $1 - h$ | $\ln w(\bar{\mathbf{z}}_{P1}, \mathbf{o})$ | $-V_{P1}$ |
| $h$ | $\ln w(\bar{\mathbf{z}}_{P2}, \mathbf{o})$ | $-V_{P2}$ |
| $p_{12}$ | $\ln w(\bar{\mathbf{z}}_{F1}, \mathbf{o}) - \frac{1}{2} (\ln w(\bar{\mathbf{z}}_{P1}, \mathbf{o}) + \ln w(\bar{\mathbf{z}}_{P2}, \mathbf{o}))$ | $-V_{F1} + \frac{1}{2} (V_{P1} + V_{P2})$ |
| $4h(1 - h) - p_{12}$ | $m(\mathbf{A}, \mathbf{A})$ | $-M(\mathbf{A}, \mathbf{A})$ |
| $p_{12}(1 - p_{12})$ | $m(\mathbf{\Delta}, \mathbf{\Delta})$ | $-M(\mathbf{\Delta}, \mathbf{\Delta})$ |
| $2p_{12}(1 - 2h)$ | $m(\mathbf{A}, \mathbf{\Delta})$ | $-M(\mathbf{A}, \mathbf{\Delta})$ |

Now, let us note that, given the quadratic fitness function of eq. 1, the mean fitness of individuals in parental population P1 is given by 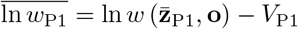. As such, we can combine the terms in each row of Table 1, to yield:

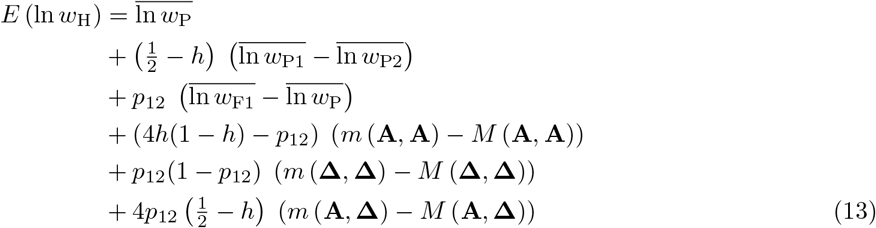

Here the overbars denote the expected fitness of randomly chosen individuals, either from a single population (subscripts P1, P2 or F1) or from the two parental populations at random (subscript P, such that 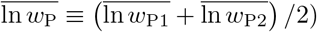.

Note that the first three terms of Equation 13 all depend on the current position of the environmental optimum, and so they capture the extrinsic or environment-dependent component of hybrid fitness. These terms depend solely on the mean log fitnesses of parental and F1 populations. By contrast, the second three terms depend only on the **A** and **Δ** – i.e. on the genomic differences accrued by the parental populations, but not on the current position of the environmental optimum. As such, these three terms capture the intrinsic, or environment-independent component of hybrid fitness. Note that fixed differences and shared polymorphisms contribute in identical ways, as long as the **A** and **Δ** are correctly defined (eqs. 3 and 6).

We note that the partition of the term shown in eq. 13 is not unique, because it includes the within-population trait variances within the extrinsic terms (Table 1). However, eq. 13 does correspond closely to the partition of Hill (1982), showing that all of the terms, including the quantities *M*(·, ·) – *m*(·, ·) are estimable as composite effects by standard quantitative genetic methods (Lande, 1981; Lynch, 1991; Lynch and Walsh, 1998, Ch. 9; Rundle and Whitlock, 2001; Schneemann et al., 2020; Clo et al., 2021). Moreover, even the separate contributions of the trait means and variances, i.e. the separate functions *M*(·, ·) and *m*(·, ·), are estimable under some conditions. This is clearest if the dominance effects are negligible (see Schneemann et al., 2022 for a discussion). In that case, all terms containing the A vanish, and the F1 trait means and variances are equal to the midparental values. As a result, Table 1 simplifies to Table 2, implying that *M*(**A**, **A**) and *m*(**A**, **A**) can be separately estimated.

**Table 2:** Components of expected log hybrid fitness with additive phenotypes

| Coefficient | $-\sum_{j=1}^n E^2(z_H - o)$ | $-\sum_{j=1}^n \text{Var}(z_H)$ |
| --- | --- | --- |
| $1 - h$ | $\ln w(\bar{\mathbf{z}}_{P1}, \mathbf{o})$ | $-V_{P1}$ |
| $h$ | $\ln w(\bar{\mathbf{z}}_{P2}, \mathbf{o})$ | $-V_{P2}$ |
| $p_{12}$ | 0 | $M(\mathbf{A}, \mathbf{A})$ |
| $4h(1 - h)$ | $m(\mathbf{A}, \mathbf{A})$ | $-M(\mathbf{A}, \mathbf{A})$ |

Even when dominance effects are non-negligible, some of the individual function values can be estimated, if fitness measurements are made in environments to which the parental populations are well adapted (Rundle and Whitlock, 2001). For example, if the mean phenotype of P1 is optimal 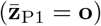, then from Table 1 and eqs. 1, 3 and 11, the log fitness of the mean P2 phenotype is 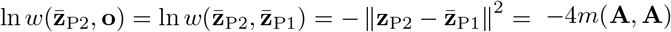. A set of equivalent results for population mean log fitness is shown in Table 3.

**Table 3:**
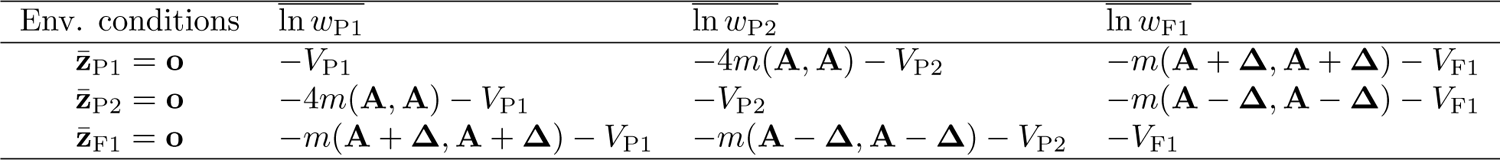
Population mean log fitnesses in different environmental conditions

| Env. conditions | $\ln w_{P1}$ | $\ln w_{P2}$ | $\ln w_{F1}$ |
| --- | --- | --- | --- |
| $\bar{\mathbf{z}}_{P1} = \mathbf{o}$ | $-V_{P1}$ | $-4m(\mathbf{A}, \mathbf{A}) - V_{P2}$ | $-m(\mathbf{A} + \Delta, \mathbf{A} + \Delta) - V_{F1}$ |
| $\bar{\mathbf{z}}_{P2} = \mathbf{o}$ | $-4m(\mathbf{A}, \mathbf{A}) - V_{P1}$ | $-V_{P2}$ | $-m(\mathbf{A} - \Delta, \mathbf{A} - \Delta) - V_{F1}$ |
| $\bar{\mathbf{z}}_{F1} = \mathbf{o}$ | $-m(\mathbf{A} + \Delta, \mathbf{A} + \Delta) - V_{P1}$ | $-m(\mathbf{A} - \Delta, \mathbf{A} - \Delta) - V_{P2}$ | $-V_{F1}$ |

If we also note the following identities:

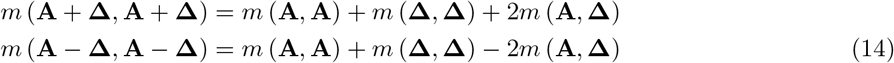

then it follows that the quantities *m*(**A**, **A**) and *m*(**A**, **Δ**) can be estimated from reciprocal transplant experiments in habitats to which the parental populations are well adapted (i.e. habitats where 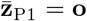 and 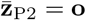). Moreover, the remaining function, *m*(**Δ**, **Δ**) can be estimated either with genetically homogeneous parental lines (i.e., if *V*_P1_ = *V*_P2_ = *V*_F1_ = 0), or with data from a third environment in which the F1 shows bounded hybrid advantage such that 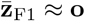.

### Interpreting the functions *m*(·, ·) and *M*(·, ·)

In the previous section, we saw that genomic differences between populations influence the mean log fitness of their hybrids solely via the functions *m*(·, ·) and *M*(·, ·), as applied to the additive and dominance effects (**A** and **Δ**). We also saw that the value of these functions can, in principle, be estimated from hybrid fitness data. In this section we show that these functions have a simple interpretation, which can be related to the divergence history of the populations.

It follows from eqs. 11 and 12, that *m*(·, ·) and *M*(·, ·) can be interpreted on a trait-by-trait basis, as the sum over the means and variances of the changes on each trait. However, it can also be helpful to consider the overall size of changes in multi-dimensional trait space, i.e. the arrows depicted in Figure 1.

To see this, let us begin by noting that the function *m*(·, ·) captures the *net effect of evolutionary change*. For example, for the additive effects, from eqs. 7 and 11 we find:

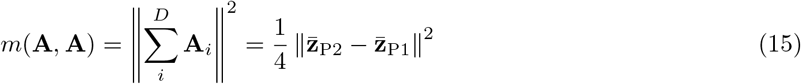

so that *m*(**A**, **A**) will be large if the evolutionary divergence between P1 and P2 led to their evolving very different phenotypes. By contrast, *m*(**A**, **A**) will be small if, due to compensatory changes at different loci, the evolutionary divergence led to little net change in phenotype. Analogous arguments apply to the dominance effects, where, from eqs. 8 and 11, the function *m*(**Δ**, **Δ**) describes the distance between the F1 and midparental phenotypes.

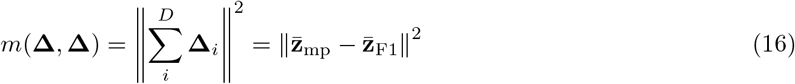

Finally, for the interaction term, we use eq. 14 from which it follows that

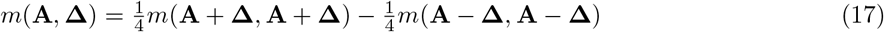

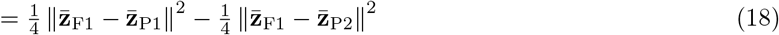

The interaction term can therefore be negative or positive, and it tells us whether the net effect of the evolutionary change has led to the F1 more closely resembling one or other of the parental populations.

If the function *m*(·, ·) describes the net effect of evolutionary change, the function *M*(·, ·), describes the *total amount of evolutionary change*. For example, from eq. 12 we have:

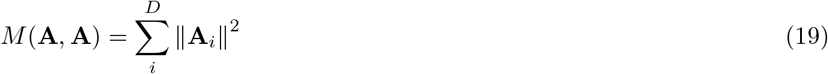

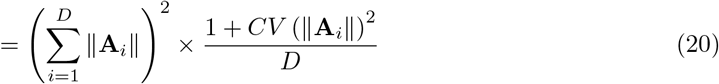

where ║**A**_*i*_║ is the length of an individual black arrow in Figure 1B, and *CV*(·) is the coefficient of variation among the complete set of *D* lengths, i.e. their standard deviation divided by their mean. It follows that *M*(**A**, **A**) will be large if there was a large amount of evolutionary change, i.e. if there were changes at many loci, and the changes were individually large. This applies regardless of whether or not the changes at each locus were compensatory, such that there was no net change in phenotype. Equation 20 also clarifies the roles of large-versus small-effect changes. It implies that for a given amount of phenotypic change (i.e. a given value of the first factor in eq. 20, or a given length of the chain of black arrows in Fig. 1B), *M*(**A**, **A**) will be larger if the changes were fewer (lower *D*) and more variable in size (higher *CV* (║**A**_*i*_║)).

All of the arguments above also apply to *M*(**Δ**, **Δ**), which concerns the chain of dominance effects; while for the interaction term, we use results analogous to eq. 14 to show that

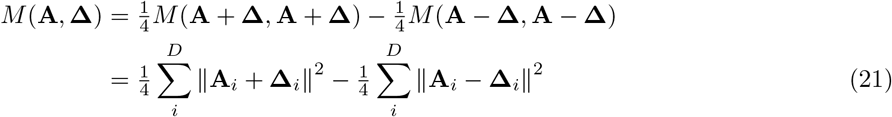

So eq. 21 will be positive if the red arrows in Figure 1A tend to be longer than the blue arrows, and vice versa. This is equivalent to asking whether the alleles that are more common in P2 tend to be phenotypically dominant. *M*(**A**, **Δ**) will be positive if P2 alleles tend to be phenotypically dominant, and negative if they tend to be phenotypically recessive.

The comments above shed light on the functions *m*(·, ·) and *M*(·, ·) individually, but eq. 13 depends on the difference between them, and this difference has its own natural interpretation. To see this, let us use eqs. 15 and 19, to show that:

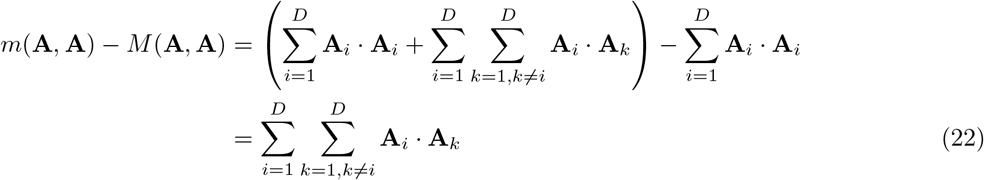

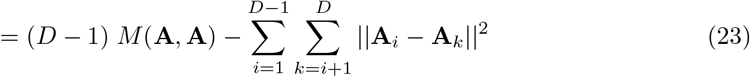

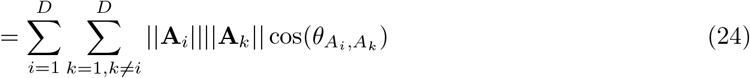

So this quantity can be interpreted in two ways. Equation 23 uses the relationship between the dot product and the squared Euclidean distance to show that *m*(**A**, **A**) – *M*(**A**, **A**) is a measure of the similarity of the evolutionary changes at different loci (Schneemann et al., 2020); it take its largest value when changes are identical at all loci (i.e. when ║**A**_i_ – **A**_*k*_║ = 0 for all *i* and *k*), but the quantity becomes smaller and negative as the effects become more different.

Similarly, eq. 24 is a generalized cosine law, and uses *θ_A_i_,A_k__* to denote the angle between the *i*th and the *k*th vectors of change (see top right of Figure 1B for an illustration). This implies that cos(*θ*) = 1 when the additive effects at two loci point in the same phenotypic direction (such that *θ* = 0); similarly, cos(*θ*) = 0 when the vectors are orthogonal (e.g., altering the values of different traits); and finally, cos(*θ*) = –1 for effects that act in opposite directions. It follows that the difference *m*(·, ·) – *M*(·, ·) quantifies the tendency for evolutionary changes at different loci to act in the same phenotypic direction. It is therefore a measure of the directionality (or conversely meandering) in the chains of evolutionary changes.

Again, the same argument applies to the chain of dominance effects (*m*(**Δ**, **Δ**) – *M*(**Δ**, **Δ**)). Finally, for the additive-by-dominance interaction, by analogy with eq. 24, we can write

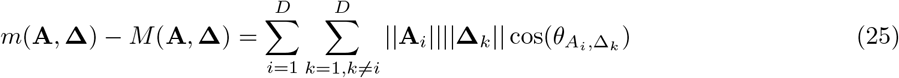

So that the interaction term measures the tendency for additive and dominance effects at different loci to point in the same phenotypic direction.

### How does directional selection affect the total amount and net effect of evolutionary change?

In the previous section we showed that the functions *m*(·, ·), *M*(·, ·) and the difference between them, *m*(·, ·) – *M*(·, ·), each have a natural interpretation. In the next two sections, we show how these quantities vary with the history of divergence between the parental lines (summarizing the results in Table 4).

**Table 4:**
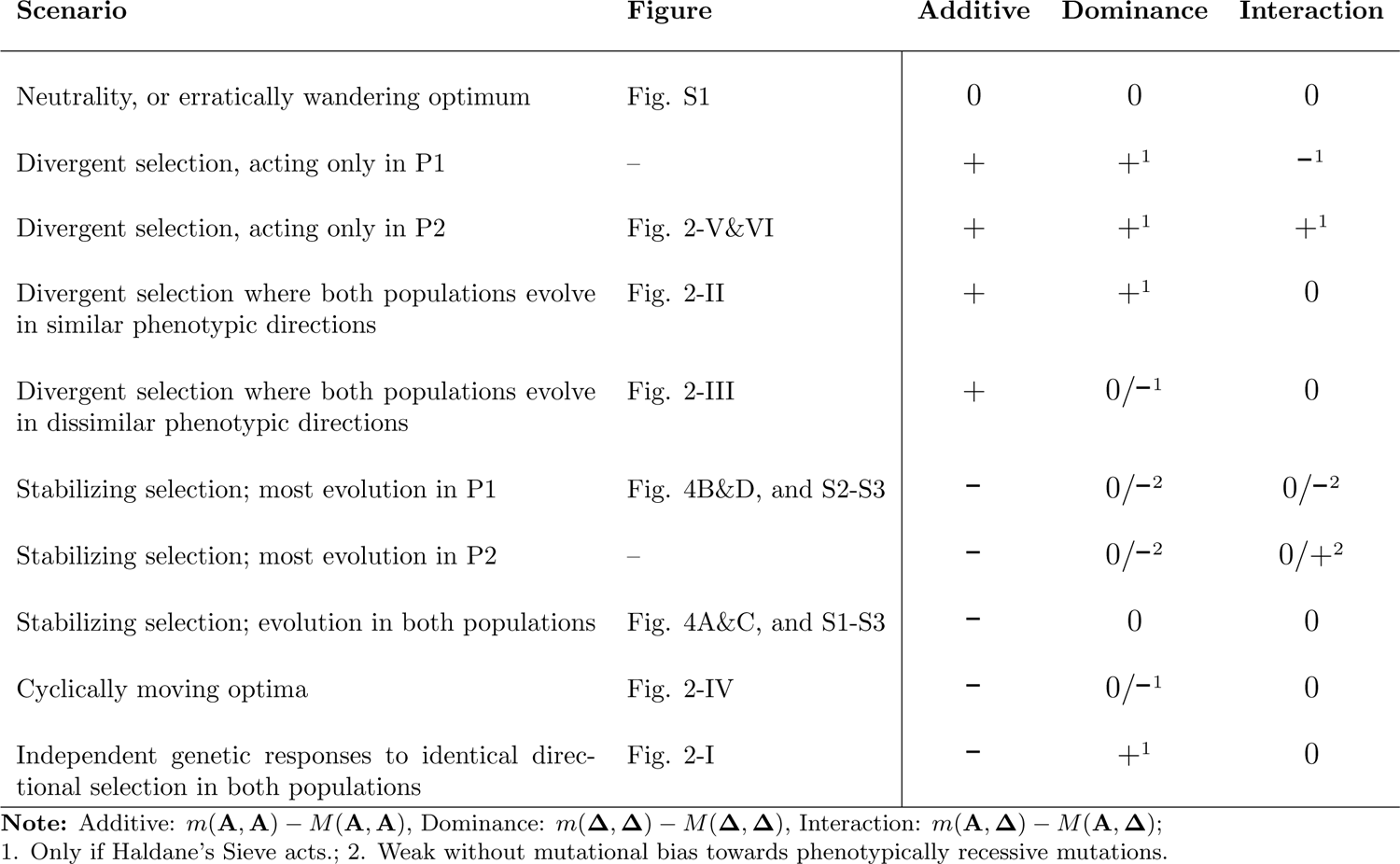
Inference of divergence scenario from the signs of terms in eq. 13

| Scenario | Figure | Additive | Dominance | Interaction |
| --- | --- | --- | --- | --- |
| Neutrality, or erratically wandering optimum | Fig. S1 | 0 | 0 | 0 |
| Divergent selection, acting only in P1 | – | + | +<sup>1</sup> | –<sup>1</sup> |
| Divergent selection, acting only in P2 | Fig. 2-V&VI | + | +<sup>1</sup> | +<sup>1</sup> |
| Divergent selection where both populations evolve in similar phenotypic directions | Fig. 2-II | + | +<sup>1</sup> | 0 |
| Divergent selection where both populations evolve in dissimilar phenotypic directions | Fig. 2-III | + | 0/–<sup>1</sup> | 0 |
| Stabilizing selection; most evolution in P1 | Fig. 4B&D, and S2-S3 | – | 0/–<sup>2</sup> | 0/–<sup>2</sup> |
| Stabilizing selection; most evolution in P2 | – | – | 0/–<sup>2</sup> | 0/+<sup>2</sup> |
| Stabilizing selection; evolution in both populations | Fig. 4A&C, and S1-S3 | – | 0 | 0 |
| Cyclically moving optima | Fig. 2-IV | – | 0/–<sup>1</sup> | 0 |
| Independent genetic responses to identical directional selection in both populations | Fig. 2-I | – | +<sup>1</sup> | 0 |

**Note:** Additive: $m(\mathbf{A}, \mathbf{A}) - M(\mathbf{A}, \mathbf{A})$ , Dominance: $m(\mathbf{\Delta}, \mathbf{\Delta}) - M(\mathbf{\Delta}, \mathbf{\Delta})$ , Interaction: $m(\mathbf{A}, \mathbf{\Delta}) - M(\mathbf{A}, \mathbf{\Delta})$ ; 1. Only if Haldane’s Sieve acts.; 2. Weak without mutational bias towards phenotypically recessive mutations.

We will begin with divergence under directional selection. To supplement verbal arguments, we use illustrative simulations of adaptive divergence under Fisher’s geometric model. Full simulation details are given in the Methods, but in brief, we used individual-based simulations, starting with a pair of identical and genetically uniform parental populations, which then evolved in allopatry to different conditions of environmental change, i.e. different positions of the phenotypic optimum (Chevin et al., 2014; Yamaguchi and Otto, 2020; Schneemann et al., 2020). While multiple variants could segregate during the simulations, the **A** and **Δ** values were calculated only for fixed differences between the populations. This means that we could avoid complications from linkage disequilibrium, which we did not treat analytically, but also implies that the analytical results apply to cases that we did not simulate.

The first set of simulations, summarized in Figure 2, involved six different divergence scenarios, illustrated by the cartoons in the left-hand panels. In scenarios I-III, both populations adapted to distant optima at a distance 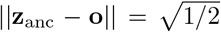 from their shared ancestral phenotype (such that their initial fitness was exp(–1/2) ≈ 60% of its maximum value). The sole difference between scenarios I-III is the relative positions of the optima experienced by each population. In scenario I, the two optima moved in identical ways, so that this scenario corresponds to mutation-order speciation (Mani and Clarke, 1990). In scenarios II-III, the two optima differed, so that these scenarios correspond to divergent selection and local adaptation (Schluter, 2000); in scenario II, the optima differed on different traits, while in scenario III, the optima differed on the same trait, but in opposite phenotypic directions. Finally, scenarios IV-VI corresponded to scenarios I-III, but with both bouts of adaptive substitution taking place in population P2, while P1 retained their common ancestral phenotype. This meant that P2 adapted to two successive changes in environmental conditions (i.e. two changes in the position of its optimum). After the initial bout of adaptation in P2, its optimum either jumped back to its initial position (scenario IV), or changed on a different trait (scenario V), or jumped again in the same phenotypic direction (scenario VI). Panels A-I of Figure 2 summarizes the results of 100 replicate simulations under each of these six scenarios, after *D* = 50 substitutions had occurred.

**Figure 2:**
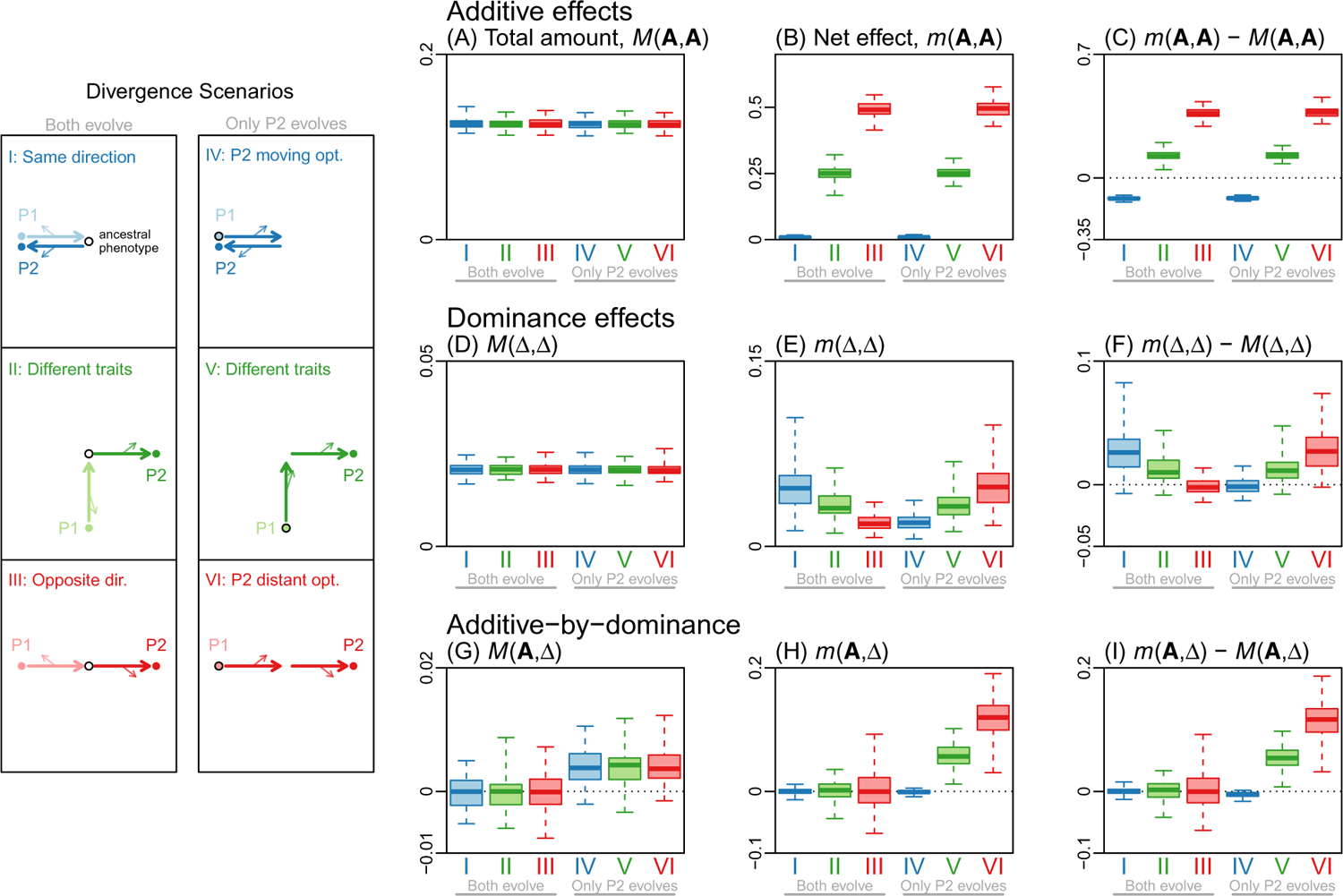
The history of directional selection affects the total amount and net effect of evolutionary change. Illustrative individual-based simulations of divergence between allopatric populations, driven by directional selection. Simulations used six distinct scenarios of divergence, illustrated via their net additive and dominance effects in the cartoons in the left-hand panels. In these panels the lighter (darker) arrows illustrate the evolutionary changes fixed in the P1 (P2) lineage. The larger arrows show the additive effects (all defined in the direction from P1 to P2), and the smaller arrows the dominance effects. The ancestral phenotype is shown by an empty black circle. Scenarios are **I**: both populations adapt to the same distant optimum; **II**: each population adapts to shifted optimum on a different phenotypic trait; **III**: each population adapts to a shifted optimum on the same trait, but in opposite phenotypic directions; **IV**: P2 alone adapts to an optimum that shifts in one phenotypic direction, and then shifts back to its initial position; **V**: P2 alone adapts to an optimum that changes on one trait, and then on another; **VI**: P2 alone adapts to an optimum that shifts twice in the same phenotypic direction. **(A)**-**(I)**: Boxes represent results for 100 replicate simulations (median, quantiles and full range), each including *n* = 20 traits, and halted after *D* = 50 fixations. The quantities shown match those in Tables 1 and 3. The quantities vary predictably between the six scenarios, and in different ways for the additive and dominance effects (see text). Simulation parameters were *N* = 1000, n = 20, and 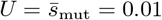.

#### Additive effects

Results for the simulated additive effects are shown in Figure 2A-C. Figure 2A shows that the total amount of evolutionary change, *M*(**A**, **A**), was identical under all six scenarios. This is because all scenarios involved two bouts of adaptive substitution under equivalent conditions; as such, they led to the same total amount of change, regardless of how the changes were distributed among the traits and the diverging populations.

Figure 2B shows the net effect of the evolutionary change, *m*(**A**, **A**). This quantity is proportional to the squared distance between the parental mean phenotypes (eq. 15). So when populations are well adapted to their optima, *m*(**A**, **A**) will be proportional to the squared distance between these optima. This explains the observed results of *m*(**A**, **A**) ≈ 0 for scenarios I and IV, *m*(**A**, **A**) ≈ 2║**z**_anc_ – **o**║^2^/4 = 0.25 for scenarios II and V, and *m*(**A**, **A**) ≈ ║2(**z**_anc_ – **o**)║^2^/4 = 0.5 for scenarios III and VI.

Figure 2C combines results from Fig. 2A-B, to quantify the directionality in the chain of additive effects that differentiate P1 and P2. From eq. 24, this value will be positive if the effects mostly point in the same direction, such that cos(*θ*) ≈ 1 holds for most pairs of changes. This occurs under scenarios III and VI, where most of the additive effects point from the P1 phenotype to the P2 phenotype. Results are also positive, but around half as large, in scenarios II and V, since cos(*θ*) ≈ 1 for half of the pairs of changes and cos(*θ*) ≈ 0 for the other half. By contrast, when natural selection tends to return the chain of additive effects to its starting point, as in scenarios I and IV, then cos(*θ*) < 0 will hold on average, leading to a negative value.

All of the quantitative results above will, of course, vary over time (as more divergence accrues), and with the various parameters of the model. For example, previous work has shown that populations often approach their optima more efficiently if the number of traits under selection, *n*, is small, because mutations tend to have fewer deleterious pleiotropic effects (e.g. Orr, 1998; Welch and Waxman, 2003; Matuszewski et al., 2014; Chevin et al., 2014). This is confirmed in Figure 3A, which shows results for scenarios II-III as a function of the divergence, D. When we reduced the number of traits from *n* = 20 to *n* = 2 populations approached their optima much more rapidly. Figure 3B shows how the relative sizes of *M*(**A**, **A**) and *m*(**A**, **A**) change with the divergence. In the initial stages of divergence, as the distant optima are approached (see Fig. 3A), the additive effects point in a consistent direction, and so the ratio decreases. More quantitatively, it follows from eq. 20 that if the changes at each locus act in the same direction, then the first term of eq. 20 will equal *m*(**A**, **A**). If these changes are also similarly sized (such that *CV*(║**A**_*i*_║) ≈ 0), then *M*(**A**, **A**)/*m*(**A**, **A**) ≈ 1/*D* should hold. This prediction - indicated by the grey line in Figure 3B - does hold approximately for scenario III when *n* = 2 (solid red line in Figure 3B), while the optimum remains distant. The decline is slower than 1/D (implying a less direct approach to the optimum), when populations fixed deleterious pleiotropic effects (*n* = 20; dashed red line), or when the position of the ancestral phenotype led to effects acting in different phenotypic directions (scenario II; green lines). The decline also slows as the optimum is approached, and populations begin to fix alleles of smaller effect (thereby increasing *CV*(║**A**_*i*_║); Orr, 1998). In all cases, the ratio *M*(**A**, **A**)/*m*(**A**, **A**) starts to increase after the optimum is reached, when evolutionary changes continue to accrue, but without much net phenotypic change (Schiffman and Ralph, 2021).

**Figure 3:**
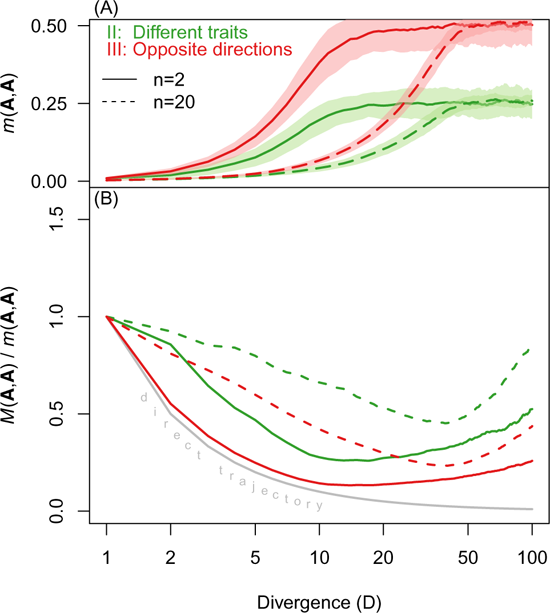
The net effect and total amount of evolution change predictably during directional selection. Panels show **(A)**: the net effect of evolutionary change in the additive effects, *m*(**A**, **A**). and **(B)**: the ratio of the total amount to the net effect, *M*(**A**, **A**)/*m*(**A**, **A**), both plotted as functions of *D*, the number of substitutions that have accumulated. Results are compared for different numbers of phenotypic traits, namely *n* = 2 (solid lines) and *n* = 20 (dashed lines), and for two scenarios detailed in Figure 2. All curves represent means over 100 replicate simulations, with shaded areas representing one standard deviation. The grey curve in **(B)** shows the prediction of *M*(**A**, **A**)/*m*(**A**, **A**) ≈ 1/D, which holds when the additive effects at each locus are identical (eq. 20). Other simulation parameters matched Figure 2 (*N* = 1000 and 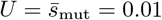).

#### Dominance and interaction terms

Results for the simulated dominance effects under the six divergence scenarios are shown in Figure 2D-F. For the total amount of evolutionary change (*M*(**Δ**, **Δ**); Fig. 2D), results are indistinguishable, just as they were for the additive effects (Fig. 2A). By contrast, results for net effect (*m*(**Δ**, **Δ**); Fig. 2E) are qualitatively different, and so - in consequence - are results in Fig. 2F.

The key fact here is Haldane’s Sieve - the tendency for directional selection to preferentially fix alleles that are dominant in the direction of past selection (Haldane, 1924, 1927; Frankham, 1990; Crnokrak and Roff, 1995; Schneemann et al., 2022), especially when adaptation takes place from new mutations, rather than standing variation (Orr and Betancourt, 2001). This means that dominance effects reflect the history of past selection in a different way to the additive effects.

The result is that for scenarios I and VI, all of the dominance effects point in a consistent direction (from the ancestral state to the new optimum); leading to large net changes in phenotype (i.e. to large *m*(**Δ**, **Δ**); Fig. 2E) and to large positive values of *m*(**Δ**, **Δ**) – *M*(**Δ**, **Δ**) (Fig. 2F). By contrast, for scenarios III and IV, the dominance effects point in opposite directions (half towards one new optimum, and half towards the other), leading to a small values of *m*(**Δ**, **Δ**) (Fig. 2D) and weakly negative values of the difference *m*(**Δ**, **Δ**) – *M*(**Δ**, **Δ**) (Fig. 2F).

Finally, results for the additive-by-dominance interactions are shown in Figure 2G-I. Unlike terms involving additive or dominance effects alone, the interaction terms tell us whether the two populations have evolved in different ways (eqs. 18, 21 and 25). As such, it is unsurprising that all of these terms are close to zero for scenarios I-III, where both populations underwent similar amounts and patterns of evolution. By contrast, for scenarios IV-VI, P2 alone adapted to a distant optima, and did so via dominant substitutions. It follows that, for these scenarios, the P2 alleles tended to be phenotypically dominant, leading to *M*(**A**, **Δ**) > 0; eq. 21; Fig. 2G). If the parental populations differ phenotypically (scenarios V-VI), then the F1 will more closely resemble the population carrying the dominant alleles (*m*(**A**, **Δ**) > 0; eq. 18; Fig. 2H). The result, shown in Figure 2I, is that the additive and dominance effects at different loci tend to point in opposite directions for scenario IV (for which *m*(**A**, **Δ**) – *M*(**A**, **Δ**) is weakly negative), but in the same phenotypic direction for scenarios V-VI (for which *m*(**A**, **Δ**) – *M*(**A**, **Δ**) is positive).

### How does stabilizing selection affect the total amount and net effect of evolutionary change?

Now let us turn to evolution under stabilizing selection. The arguments in this section are illustrated by simulation results shown in Figure 4. In these simulations, the optima for both populations remained stationary and identical, matching their common ancestral phenotype. As such, any evolutionary change was due to the drift-driven fixation of mildly deleterious mutations, combined with compensatory changes.

**Figure 4:**
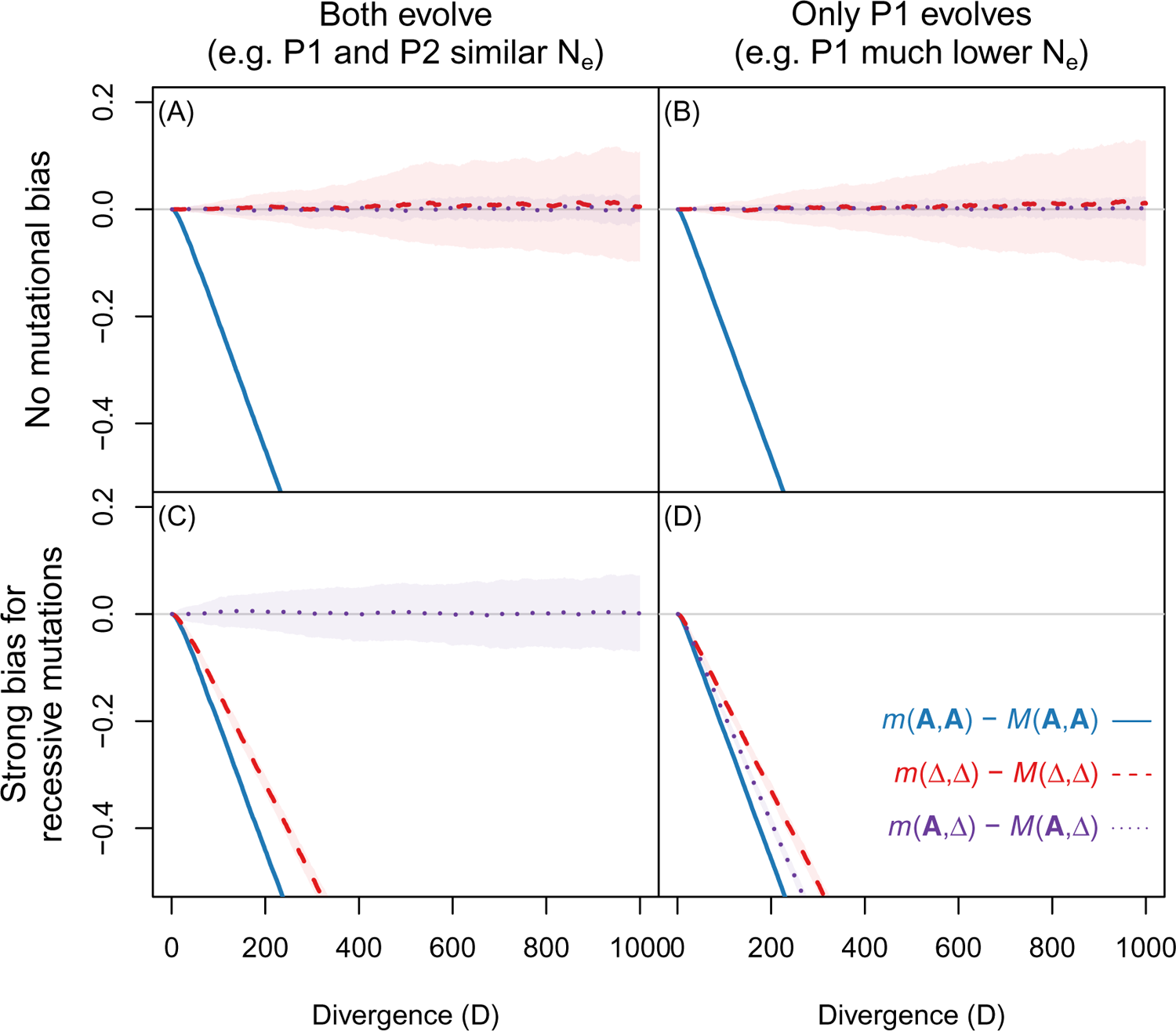
The net effect and total amount of evolution change predictably under stabilizing selection. Each plot compares the amount of directionality in the additive effects (*m*(**A**, **A**) – *M*(**A**, **A**); solid blue lines), dominance effects (*m*(**Δ**, **Δ**) – *M*(**Δ**, **Δ**); dashed red lines), and the interaction term (*m*(**A**, **Δ**) – *M*(**A**, **Δ**); dotted purple lines), plotted against the level of genetic divergence (D) under stabilizing selection to a stationary optimum. **A**-**B**: results with the standard model of mutation (as in Figure 2), with all mutations equally likely to be phenotypically recessive or dominant. **C**-**D**: results with biased mutation, in which mutations of larger phenotypic effect were more likely to be recessive (see Appendix 2). **A** and **C**: Both populations had identical population sizes of *N* = 100, so that they accrued substitutions at a similar rate; **B** and **D**: We assumed that P2 remained in the optimal ancestral state, while P1 (with *N* = 100) underwent all of the evolutionary change. Lines and shaded areas represent the mean and one standard deviation across 200 replicate simulations. Other simulation parameters matched Figure 2 (*n* = 20 and 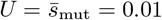).

#### Additive effects

The first key point about stabilizing selection is that the net phenotypic change, *m*(**A**, **A**), will reach a stochastic equilibrium, reflecting the deviations of the populations from the optimum due to mutation and drift. Barton (2016) showed that, with independent loci but otherwise very general assumptions, the expected log fitness under stabilizing selection on n traits is ~ –*n*/(4*N_e_*) (see also Lande, 1976; Hartl and Taubes, 1996; Poon and Otto, 2000; Zhang and Hill, 2003; Tenaillon et al., 2007; Lourenço et al., 2011; Chevin et al., 2014; Roze and Blanckaert, 2014). Now, if the two populations are maladapted in random phenotypic directions (such that their displacements from the optimum are orthogonal on average; Schneemann et al., 2022), then it follows from eqs. 1 and 15, that

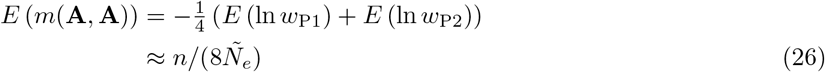

where *Ñ_e_* is the harmonic mean of the two effective population sizes. This result is confirmed by simulations reported in Appendix 2 as shown in Supplementary Figure S1.

While the net effect of change is determined largely by *n* and *N_e_*, the total amount of change will depend on the size of mutations that fix (as determined by the distribution of scaled selective effects: *N_e_s*). Evolutionary changes will continue to accrue even after *m*(**A**, **A**) has equilibrated (Schiffman and Ralph, 2021), so that *M*(**A**, **A**) will increase over time at a constant rate. The result is illustrated by the solid blue lines in Figure 4A-D, which show that *m*(**A**, **A**) – *M*(**A**, **A**) declines steadily under stabilizing selection.

#### Dominance and interaction terms

The evolution of dominance effects under stabilizing selection is more complex, and sensitive to the underlying model of mutation. For this reason, some of the discussion is relegated to Appendix 2, while here we report the clearest patterns.

Figure 4A-B show results with the mutation model used in Figure 2, in which each new mutation was equally likely to be phenotypically recessive or phenotypically dominant. In this case, we found that *m*(**Δ**, **Δ**) ≈ *M*(**Δ**, **Δ**) at all levels of divergence (dashed red lines), because *m*(**Δ**, **Δ**) and *M*(**Δ**, **Δ**) both increased with *D*, but at identical rates. The reason is that, unlike the additive effects, the dominance effects are not expressed together in the parental genotypes during the divergence process, and so unlike the additive effects, the dominance effects show little tendency to be coadapted to their optimum, but are free to wander in phenotypic space (Schneemann et al., 2020, 2022).

Figure 4C-D shows comparable results when we adopted the mutational model of Schneemann et al. (2022), in which larger effect mutations were more likely to be phenotypically recessive (Billiard et al., 2021; see Appendix 2 for full details). Now, as shown by the dashed red lines, *m*(**Δ**, **Δ**) – *M*(**Δ**, **Δ**) decreases over time. This is because both *M*(**Δ**, **Δ**) and *m*(**Δ**, **Δ**) increase with *D*, but at different rates. This implies that the dominance effects, too, have a tendency to be coadapted to the optimum. The explanation is clear if we consider the extreme case of complete phenotypic recessivity. In that case, the additive and dominance effects of mutations would be equal and opposite (such that the heterozygous effects were zero). As such, the apparent “coadaptation” of the dominance effects would follow trivially from the coadaptation of the additive effects (see Appendix 2 for more details). The dominance curves in Figure 4C-D show this effect in less extreme form, so that *m*(**Δ**, **Δ**) – *M*(**Δ**, **Δ**) decreases with *D*, but slightly less rapidly than *m*(**A**, **A**) – *M*(**A**, **A**).

Consider finally the interaction terms, shown by the dotted purple lines in Figure 4. As shown in Figure 4A and C, the interaction terms are always close to zero when both populations undergo similar patterns of evolution (in this case due to their identical population sizes). More surprisingly, as shown in Figure 4B, with the standard model of mutation, results remain qualitatively unchanged when P2 remained in its ancestral state, while all of the evolution took place in P1. The explanation is that, with this mutation model, the evolving population showed no tendency to fix phenotypically recessive mutations – and recalling that, under this model, mutations can be recessive for fitness, even if they are additive, or even dominant, for the phenotype (Manna et al., 2011). By contrast, when mutations tended to be phenotypically recessive (Figure 4C-D) then *M*(**A**, **Δ**) becomes non-zero, and the interaction term becomes a reliable guide to whether the recessive mutations were fixed more-or-less equally in both populations (such that *m*(**A**, **Δ**) ≈ *M*(**A**, **Δ**) ≈ 0; Figure 4C), or mostly in P1 (*m*(**A**, **Δ**) – *M*(**A**, **Δ**) < 0; Figure 4D) or in P2 (*m*(**A**, **Δ**) – *M*(**A**, **Δ**) > 0; not shown). Note that this signal would remain even after a transient reduction in *N_e_*, as long as a substantial number of phenotypically recessive mutations were fixed during the bottleneck.

## Discussion

This work has explored how the mode of divergence between parental populations impacts the fitness of their hybrids. We have focused on expected hybrid fitness, and not the variance or higher moments, and on results that apply to controlled crosses, where the measures of genome composition (*h* and *p*_12_) are probabilities determined by the crossing scheme. However, as we show in Appendix 1, the results can also be applied to data of other kinds, e.g. when *h* and *p*_12_ are estimates of ancestry from individual genome sequences. To generate simple, testable predictions, we have used a simple model of selection on quantitative traits introduced by Fisher (1930), but have extended and generalized previous work on this model, both by allowing for arbitrary additive and dominance effects at each locus, and by accounting for segregating variation within the parental populations.

Results show how the expected fitness of hybrids depends on only a handful of summary statistics, which describe the evolutionary changes that differentiate the populations, and which are described by the functions *m*(·, ·) and *M*(·, ·) (eqs. 11–12). If the population genetic parameters, or the history of environmental change, influence the outcomes of hybridization (Chevin et al., 2014; Yamaguchi and Otto, 2020; Schneemann et al., 2020), then they do so via these quantities. The statistics, moreover, are estimable by quantitative genetic methods (Hill, 1982; Lynch, 1991; Rundle and Whitlock, 2001; Schneemann et al., 2020; Clo et al., 2021), and have a natural interpretation. In particular, m(·, ·) represents the “net effect of evolutionary change”, *M*(·, ·) represents the “total amount of evolutionary change”, and the difference *m*(·, ·) – *M*(·, ·) (which appears directly in eq. 13) represents the similarity of changes at different loci (eqs. 24–25; Martin et al., 2007; Chevin et al., 2014; Fraïsse and Welch, 2019). Applied to additive effects, *m*(**A**, **A**) – *M*(**A**, **A**), closely resembles an *Q_ST_-F_ST_* comparison (Whitlock, 2008).

It follows immediately from the results above that very different histories of evolutionary divergence can yield identical patterns of hybrid fitness, as long as they lead to the same values of *m*(·, ·) – *M*(·, ·). Nevertheless, we have shown that some information about the divergence history is present in hybrid fitness data (Figure 2). These results are summarized in Table 4, which contains the predicted signs of the key quantities that appear in the three final terms in eq. 13.

As is clear from Table 4, the simplest results concern directional selection. In particular, *m*(**A**, **A**) – *M*(**A**, **A**) will tend to be positive only when the divergence between the parental lines was driven by positive selection towards distinct environmental optima. The size of the term will depend on further details of the adaptive divergence (Figure 3). It is maximized, for example, when all allelic changes produced identical effects (eq. 23), and decreases in size if the adaptive change is achieved via a circuitous route (e.g. because of deleterious pleiotropy, overshoots of the optimum, fluctuating environmental conditions, or maladapted ancestral states); and – for a given amount of phenotypic change - the term decreases if the number of loci is smaller, and their effects more variable in size (eq. 20; see also Chevin et al., 2014). Additional and complementary information about the divergence history is present in the dominance and interaction terms (*m*(**Δ**, **Δ**) – *M*(**Δ**, **Δ**) and *m*(**A**, **Δ**) – *M*(**A**, **Δ**)). Due to Haldane’s Sieve (Haldane, 1924), dominance effects will often point in the direction of past selection. For example, if one population adapted to new conditions via dominant mutations, while the other remained in their shared ancestral habitat, then we would expected both *m*(**Δ**, **Δ**) –*M*(**Δ**, **Δ**) and *m*(**A**, **Δ**) –*M*(**A**, **Δ**) to be positive, as well as m(**A**, **A**) –*M*(**A**, **A**). It follows, therefore, that the analysis of hybrid fitness might tell us not only about the presence of past directional selection (e.g. Fraser, 2020), but also about the direction of that selection, and the lineage in which the adaptation occurred (see Figure 2; Table 4).

If *m*(**A**, **A**)–*M*(**A**, **A**) is negative, then inferences about the evolutionary divergence are more challenging, since negative values can arise in a number of different ways (see Figures 2 and 4 and Table 4). Nevertheless, even in this case, the dominance and interaction terms might yield useful information. Consider, for example, a pair of populations with similar current phenotypes and fitness, but which nonetheless produce unfit hybrids, due to *m*(**A**, **A**) – *M*(**A**, **A**) ≪ 0. In this case, an estimate of *m*(**Δ**, **Δ**) – *M*(**Δ**, **Δ**) ≈ 0 would not be very informative, as it can arise under stabilizing selection, fluctuating selection, or even directional selection if Haldane’s Sieve is weak (Orr and Betancourt, 2001). However, a strongly positive estimate of m (**Δ**, **Δ**) – *M*(**Δ**, **Δ**) would be consistent with the populations having diverged via different genomic responses to identical directional selection (Figure 2 scenario I). By contrast, if this dominance term were negative, and the interaction term was also non-zero, then this would be consistent with one of the populations having undergone prolonged periods of low *N_e_*, and fixing deleterious recessive mutations (Figure 4D). The sign of the interaction term, *m*(**A**, **Δ**) – *M*(**A**, **Δ**), would then tell us which of the two populations had experienced the low *N_e_*. Note that, from eq. 13 the result would be alleles from one parental line being selected against, despite the lines having equal fitness (Barton, 1992).

A major caveat of all of the results presented here is the extreme simplicity of the phenotypic model (with its lack, for example, of phenotypic epistasis, and directional plasticity; Stamp and Hadfield, 2020). However, this model can be defended as an approximation of more complex and realistic models (Martin, 2014), or simply as a way of generating a fitness landscape with few parameters (Simon et al., 2018). In this case, as shown in Appendix 1, we can follow Chevin et al. (2014), and reframe our results in terms of fitness effects, rather than phenotypic changes. Of course, even as a fitness landscape, the quadratic model of eq. 1 remains very simple, and precludes strong fitness epistasis and multi-locus fitness interactions (Barton, 2001; Martin et al., 2007; Fraïsse and Welch, 2019) - both of which are often observed in cross data (Coyne and Orr, 2004; Fraïsse et al., 2014, 2016). Yet even in the presence of such effects, results might still apply to transformed fitness measurements (Fraïsse et al., 2016; Simon et al., 2018; Schneemann et al., 2020).

A second major caveat is our neglect of linkage disequilibrium (Lande, 1981; Schneemann et al., 2020), which is essential to studying the full dynamics of introgression. Nevertheless, even the current results have suggestive implications for the stability of local adaptation, and the evolution of genetic architectures (Dekens et al., 2021; Yeaman, 2022). For example, the dominance of alleles may be a major determinant of the effective rates of migration between demes, and the possibility of allele swamping (Barton, 1992). Directional dominance, resulting from local adaptation, may therefore act as a source of asymmetric gene flow between derived and ancestral populations. Similarly, a body of previous work suggests that the architecture of adaptation will be affected by the presence or absence of gene flow (as reviewed in Yeaman, 2022). In particular, adaptation in the face of gene flow should create architectures that are more “concentrated”, i.e., involving fewer, larger effects, and tighter linkage. Combined with results here (eq. 20), this implies that ongoing gene flow during local adaptation might sometimes increase the strength of resulting intrinsic RI.

## Methods

### Derivation of main result

We assume that individuals from our two diploid parental populations, P1 and P2, vary at *D* biallelic loci. We can arbitrarily choose one allele at each locus to be the focal allele, denoted B, such that the other allele can be denoted b. Since loci are assumed to be independent, let us first specify the genetic model for a single locus, following the standard conventions of quantitative genetics (e.g. Lynch and Walsh, 1998, Ch. 4). Accordingly, we define the contribution of the bb genotype to the trait *j* as 0, so that the point (0, 0,…, 0) in *n*-dimensional trait space corresponds to the individual with only bb genotypes at each of the *D* loci. The contribution of the Bb genotype on locus *i* to the trait *j* is defined as *a_ij_* + *d_ij_*, and the contribution of the BB genotype on locus *i* to trait *j* is 2*a_ij_*. This is summarized in Table 5.

**Table 5:** The genotypic values for locus *i* and trait *j*

| Locus $i$ genotype | Contribution to trait $j$ |
| --- | --- |
| bb | 0 |
| Bb | $a_{ij} + d_{ij}$ |
| BB | $2a_{ij}$ |

#### Properties of the three focal populations

Here we will specify properties of three key populations, namely the two parental populations, P1 and P2, and the initial F1 cross. Crucially, these populations correspond to the three possible ancestry states of any given locus in the hybrid, i.e. either both alleles are derived from P1, or both from P2, or there is mixed ancestry with one allele derived from each population. Table 6 gives a list of fundamental parameters in our model in each of these three populations.

**Table 6:**
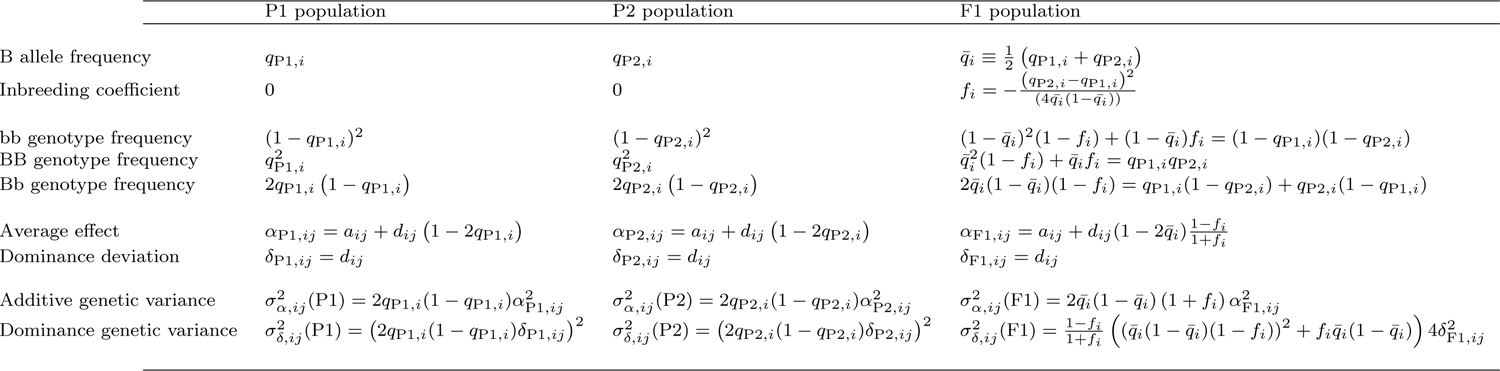
Fundamental parameters in the three focal populations at locus *i* and trait *j*

Table 6 begins by defining the marginal frequency of the focal (B) allele at locus *i* as *q*_*P*1,*i*_ and *q*_*P*2,*i*_ in populations P1 and P2 respectively. The marginal frequency of the B allele in the F1 population is the mean of the marginal frequencies in P1 and P2, denoted 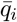. By assumption, the two parental populations are at Hardy-Weinberg equilibrium, but the F1 population will have an excess of heterozygotes, which can be parameterized by a negative coefficient of inbreeding, *f_i_*. The frequencies of the three possible genotypes at the locus, bb, Bb and BB, then follow from standard results (e.g., Lynch and Walsh, 1998, eqs. 4.21). The F1 genotype frequencies can also be written in terms of the parental allele frequencies (for example, the F1 bb frequency is the product of the marginal frequencies of the b allele in P1 and P2), which allows us to solve for the inbreeding coefficient, as shown in the Table. The next lines of the Table follow standard quantitative genetics (e.g. Fisher, 1930; Cockerham, 1954; Lynch and Walsh, 1998, Ch. 4) and define the average effects and dominance deviations of an allelic substitution at the locus in each of the populations (see, e.g., eqs. 4.10b and 4.22 in Lynch and Walsh, 1998).

These are all of the results needed to derive eqs. 3–6. Let us begin with the contribution to the mean of trait *j* from locus *i* in populations P1 and P2. This is given by the sum of the three genotype frequencies in the population, weighted by their trait contributions, as given in Table 5.

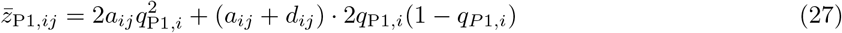

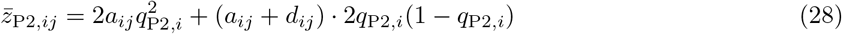

in populations P1 and P2 respectively. Equation 3 then follows immediately as

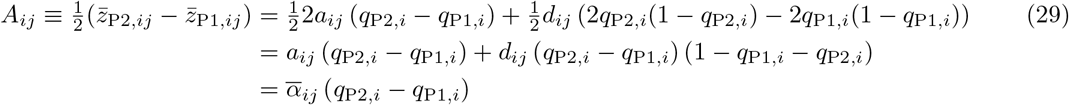

where the mean average effect is defined as

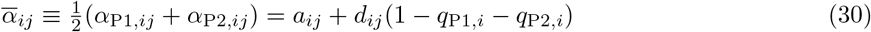

Similarly, to derive eq. 6, we use the genotype frequencies for the F1 as shown in Table 6, to yield the contribution of locus *i* to the mean of trait *j* in the F1

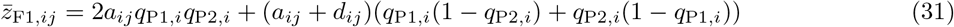

and so it follows that

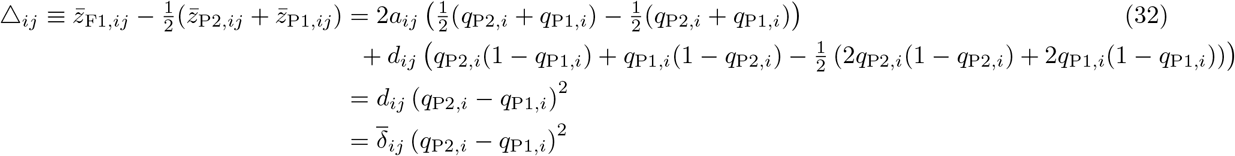

which is equation 6, and where the mean dominance deviation is simply

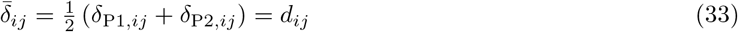

Having defined the mean trait values of each population, let us now consider their variances. The contribution of locus *i* to the variance in trait *j* in population P1 is

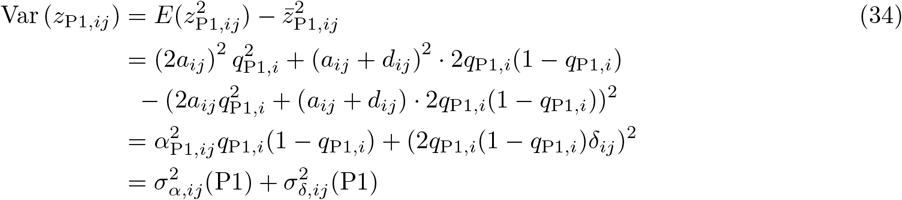

where we have partitioned the result into an additive variance and a dominance variance term, as listed in Table 6, and following eqs. 4.12 of Lynch and Walsh (1998). Similarly for P2,

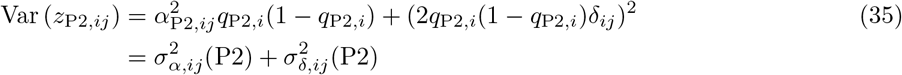

and for the F1

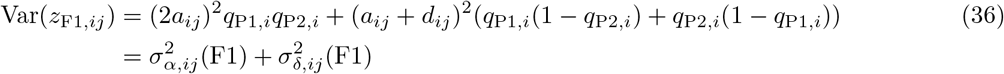

which all agree with results in Cockerham (1954). So far, we have given the contributions of a single locus to a single trait. The general results, found in Table 1, simply require summing over all loci *i* = 1,…, *D* and all traits *j* = 1,…, *n*. That is, we can write the sums of trait variances for P1, P2 and F1 as

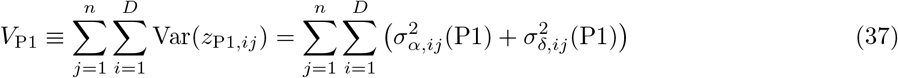

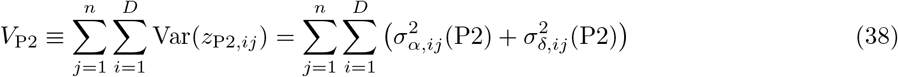

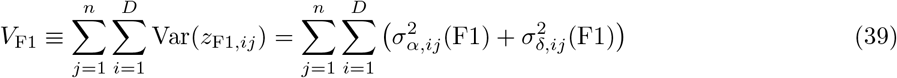

#### Extension to an arbitrary hybrid

Now, to derive the results found in Table 1 and eq. 13, let us consider an arbitrary hybrid. Let us begin by parameterizing the hybrid’s genome using the probabilities *p*_1_, *p*_2_ and *p*_12_, which are the probabilities that a randomly chosen locus in the hybrid is in each of the three possible ancestry states. That is, *p*_1_ is the probability that a randomly chosen locus in the hybrid inherits both alleles from the P1 population, *p*_2_ that it inherits both alleles from the P2 population, and *p*_12_ that it inherits one allele from each population (as with all loci in the F1). It therefore follows that

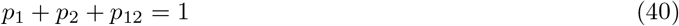

We also define the hybrid index

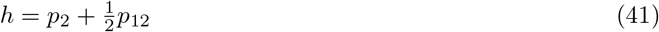

as the probability that a randomly chosen single allele in the hybrid has P2 ancestry.

Using results in Table 6, it then follows that the probabilities of the BB and Bb genotypes at a locus *i* in the hybrid are

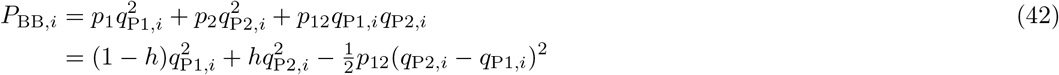

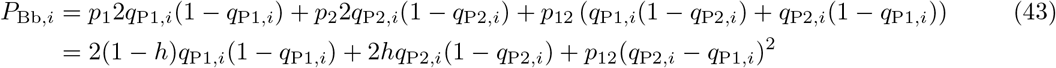

so the overall marginal probability of the B allele is

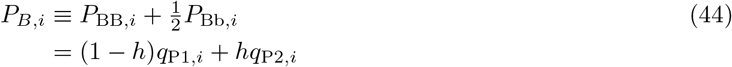

We can now derive Equation 13. First, the contribution to the mean trait value for the hybrid at locus *i* and trait *j* is

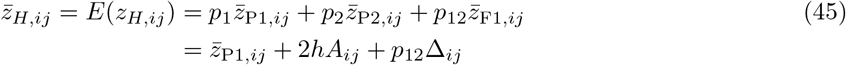

which can be seen by substituting in equations 29 and 32. Summed over the *D* loci, we have

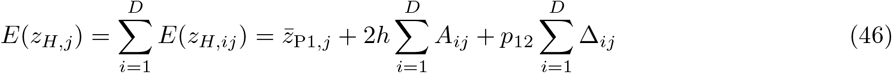

Let us now compute *E*(*z_H,j_* – *o_j_*)^2^, which appears in the first term of eq. 9. It will first be useful to define the intermediate variable

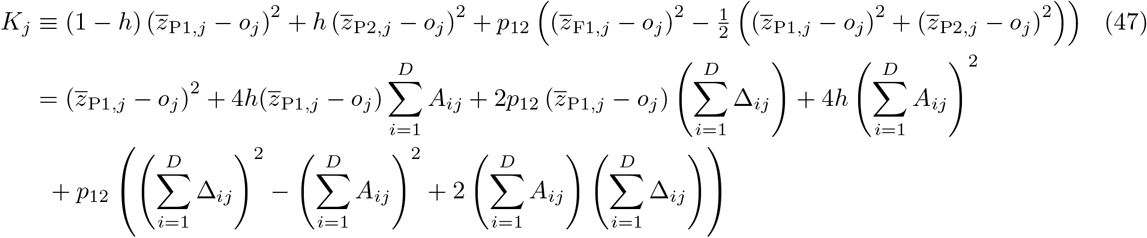

such that

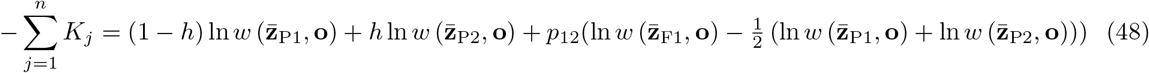

which corresponds to the sum of the top three rows for the squared mean term in Table 1.

Then we find by Equation 46,

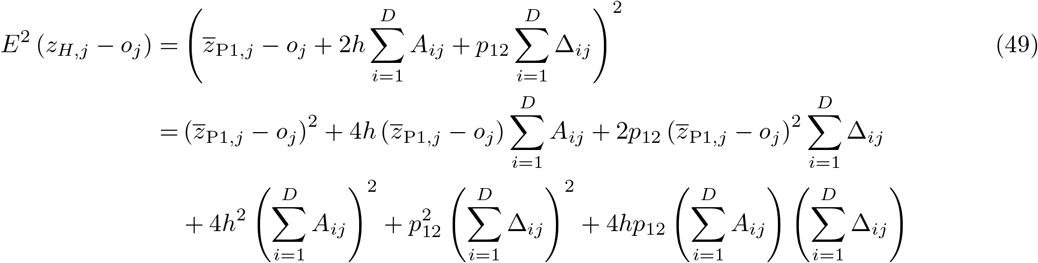

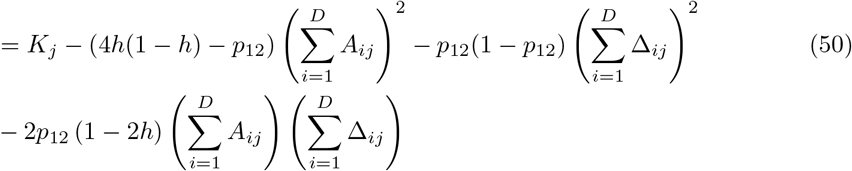

Summing over traits and using the definition of the function *m*(·, ·) in eq. 11, we can see that

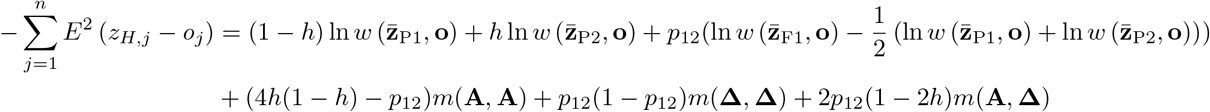

as given in the second column of Table 1.

The calculation for the variance follows in the same way, but is much more involved algebraically. The result, as shown in the third column of Table 1, is

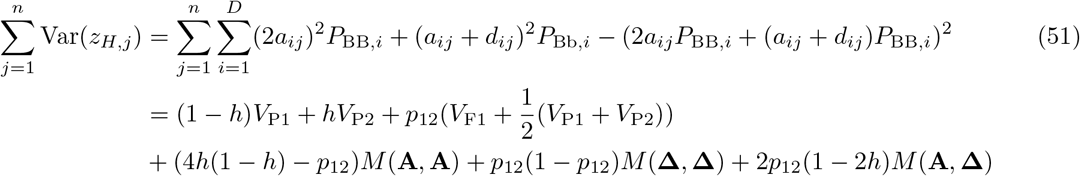

where *V*_P1_, *V*_P2_ and *V*_F1_ are defined as in eqs. 34–36, and the function *M*(·, ·) is defined by eq. 12. The first equality follows from the definition of variance and the independence of loci. The second follows by substituting variables as per their definitions above. Because the full proof is rather lengthy, although straightforward, we provide a proof in the form of a Mathematica notebook instead of writing it out here, available at https://github.com/bdesanctis/mode-of-divergence.

### Simulations

The illustrative simulations shown in Figures 2–4, calculated new quantities from runs reported previously by Schneemann et al. (2022) (and which were themselves based on the simulation methods reported in Schneemann et al., 2020). Simulations were individual-based, and used pairs of allopatric (i.e. independently simulated) populations. The populations followed the Wright-Fisher assumptions, and contained *N* simultaneous hermaphrodites, with discrete non-overlapping generations. Every generation, parents were selected with a probability proportional to their fitness (as calculated from eq. 1) with n traits under selection. Gametes were generated from the parental genomes with free recombination among all sites, and mutation. For mutation, a Poisson-distributed number, with mean 2*NU*, of mutations were randomly assigned to unique sites, and we set *U* = 0.01. The n homozygous effects for each new mutation were drawn from a multivariate normal distribution with zero mean and no covariances, and a common variance set such that the mean deleterious effects of a mutation in an optimal background was 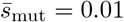. The heterozygous effect of each mutation on each trait was set at its homozygous effect multiplied by a beta-distributed random number, with bounds at 0 and 1 (corresponding to complete recessivity or complete dominance), a mean *μ* = 1/2 (implying additivity on average), and a variance of *ν* = 1/24 (Schneemann et al., 2022). After a total of D substitutions had fixed across both populations, the two parental genotypes were chosen as the genotypes containing only the fixed effects in each population. For Figures 2–3 one or both populations adapted to a optimum at a distance 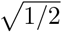 from its ancestral phenotype. In scenarios I-III, both populations in this way, while for scenarios IV-VI, we re-analysed the same simulations, but we treated all substitutions as if they had occurred in P2 while P1 remained in their common ancestral state. This was done by the contrivance of combining the first 25 substitutions accrued in two simulated populations, ensuring, therefore, that the total amount of evolutionary change was identical across all six scenarios.

## Appendix 1: Results with homogeneous parental populations

In this Appendix, we show (1) how our results apply to data where the ancestry proportions of the hybrid genome are known, and (2) how results can be expressed in terms of selective effects, rather than phenotypic changes. In both cases, for reasons explained below, we will rely on the additional assumption that parental populations are genetically homogeneous. In particular, we will assume that the focal B allele is fixed in P2 but absent in P1, such that all *q*_P2*i*_ = (1 – *q*_P1*i*_) = 1. It therefore follows from eqs. 29 and 32 that the between-population differences at each locus (eqs. 7–8) correspond directly to the genotypic effects at that locus (Table 5) i.e.

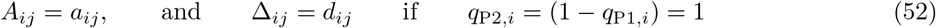

It will also be useful to rearrange the results shown in Table 1 so that they are expressed in terms of the three probabilities *p*_1_, *p*_2_ and *p*_12_ rather than the two probabilities *h* and *p*_12_ (see eqs. 40–41). Accordingly, using eqs. 11–12 and 40–41, and substituting in eq. 52 to account for the genetic homogeneity of the parental lines, we have the result shown in Table S1.

**Table S1:** Components of log hybrid fitness with homogeneous parental populations

| Coefficient | $-\sum_{j=1}^n E^2(z_H - o)$ | $-\sum_{j=1}^n \text{Var}(z_H)$ |
| --- | --- | --- |
| $p_1$ | $\ln w(\mathbf{z}_{P1}, \mathbf{o})$ | 0 |
| $p_2$ | $\ln w(\mathbf{z}_{P2}, \mathbf{o})$ | 0 |
| $p_{12}$ | $\ln w(\mathbf{z}_{F1}, \mathbf{o})$ | 0 |
| $p_1 p_{12}$ | $m(\mathbf{a} + \mathbf{d}, \mathbf{a} + \mathbf{d})$ | $-M(\mathbf{a} + \mathbf{d}, \mathbf{a} + \mathbf{d})$ |
| $p_2 p_{12}$ | $m(\mathbf{a} - \mathbf{d}, \mathbf{a} - \mathbf{d})$ | $-M(\mathbf{a} - \mathbf{d}, \mathbf{a} - \mathbf{d})$ |
| $p_1 p_2$ | $m(2\mathbf{a}, 2\mathbf{a})$ | $-M(2\mathbf{a}, 2\mathbf{a})$ |

Note that with homogenous populations, $p_1$ , $p_2$ and $p_{12}$ are now the probabilities of the three genotypes, bb, BB and Bb, as well as the ancestry states. Moreover, the arguments of the functions $M(\cdot, \cdot)$ and $m(\cdot, \cdot)$ now correspond to the phenotypic effects of inserting single alleles in either heterozygous or homozygous state into a fixed background.

Note that with homogenous populations, *p*_1_, *p*_2_ and *p*_12_ are now the probabilities of the three genotypes, bb, BB and Bb, as well as the ancestry states. Moreover, the arguments of the functions *M*(·, ·) and *m*(·, ·) now correspond to the phenotypic effects of inserting single alleles in either heterozygous or homozygous state into a fixed background.

### Results with known ancestry proportions

In the main text, we treated the quantities *h* and *p*_12_ (or equivalently, *p*_1_, *p*_2_ and *p*_12_) as probabilities determined by the crossing scheme. However, for some data, the ancestries of hybrids can be estimated directly from genome sequences. Moreover, if the parental populations are genetically homogeneous (as assumed in Table S1), then the ancestry proportions for divergent sites can be known with certainty. In this section, we show that our results also hold approximately for such data.

If *p*_1_, *p*_2_ and *p*_12_ are known proportions, instead of probabilities, loci in the hybrid become non-independent, but in a simple way so that results can be derived with basic combinatorics. For example, given some *D,p*_12_ and *p*_2_, we can choose any *Dp*_12_ out of *D* sites to be heterozygous, and any *Dp*_2_ out of the remaining *D*(1 – *p*_12_) sites to be homozygous for the allele from the second parental population, so there will be a total of

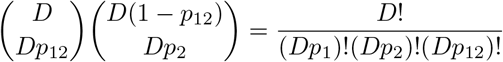

possible hybrids, and by assumption, each has equal probability. In theory, one could write out the complete discrete probability distribution function for the hybrid fitness over all possible hybrids in a given situation. One can also compute arbitrary moments using the same indicator function approach as detailed below (see also Chevin et al., 2014).

To calculate expected hybrid fitness, let *J*_1_ be the subset of the *D* loci in the hybrid that are homozygous for the P1 allele, *J*_2_ be the subset of the loci that are homozygous for the P2 allele, and *J*_12_ the subset of loci that are heterozygous. The sizes of these sets are then:

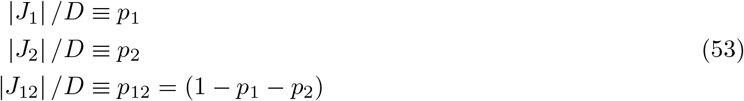

Since all divergent loci must be in one of these three states, any two of these sets can completely characterize the hybrid. We can therefore write the *j*-th trait value of an arbitrary hybrid as:

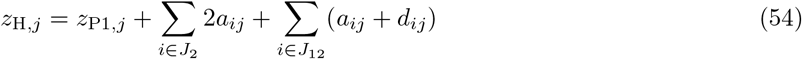

Let us now drop the subscript *j* for brevity, and calculate the expected squared deviation of the trait value from its optimum:

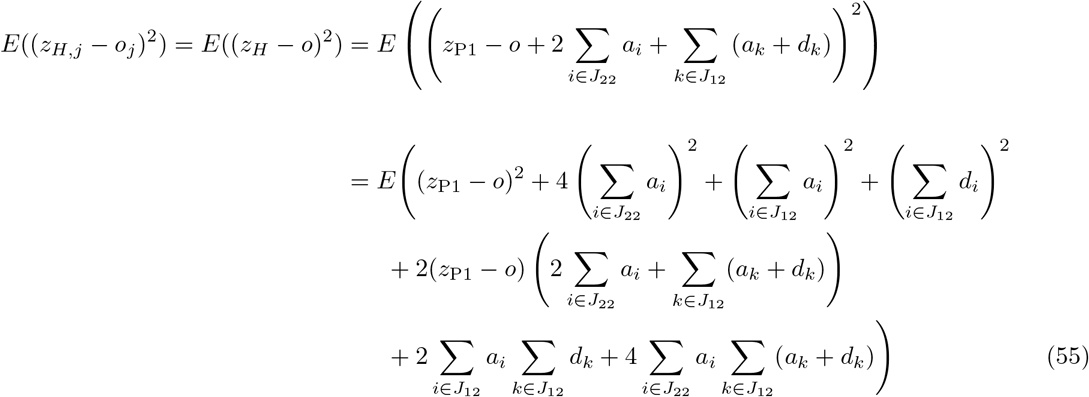

In these expressions, the expectations are not over the additive and dominance effects, but over the particular set of loci that are homozygous and heterozygous in the hybrid. That is, they are over the sets *J*_22_ and *J*_12_. To obtain expectations over these sets, we define indicator functions.

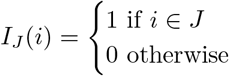

Using *x* and *y* as placeholder variables, we can then use these functions as follows:

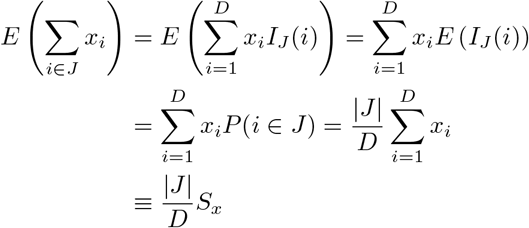

where |*J*| is the size of the set. We have introduced the notation

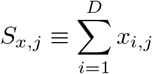

Let us also introduce

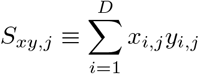

For both, we will again leave out the subscript *j* for brevity.

For the square and cross-terms in eq. 55, we use the same approach.

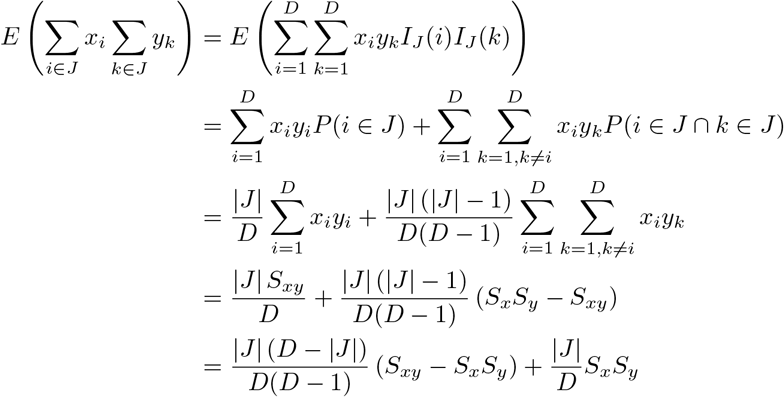

and similarly

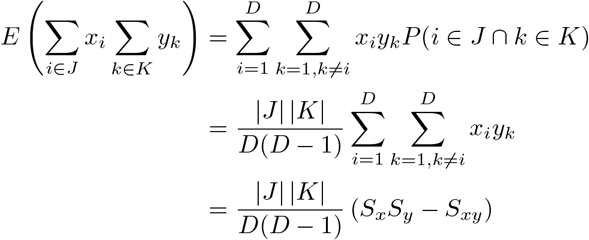

Now we can combine these results, with eqs. 53 and 55. After some algebra, we obtain

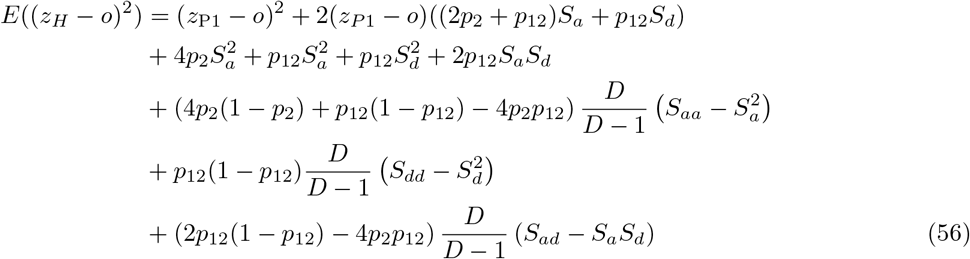

Some rearranging, and summation over traits, yields

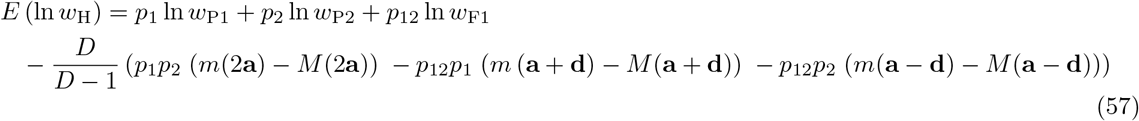

The sole difference between eq. 57 and the results summarized in Table S1 is that the functions *m*(·, ·) and *M*(·, ·) are now weighted by a new factor *D*/(*D* – 1) – which stems from the non-independence among loci when true ancestry proportions are known. Note too that *D*/(*D* – 1) ≈ 1 when the number of divergent sites is large. It follows, therefore, that the results in the main text apply approximately to data with known ancestry proportions.

### Results in terms of selective effects

We will now follow Chevin et al. (2014) and show how results can be expressed in terms of the fitness effects of alleles, rather than their phenotypic effects. This implies that the quantities *M*(·, ·) and *m*(·, ·), which describe the total amount and net effect of evolutionary change, may have a simple interpretation, even when the phenotypic model cannot be interpreted literally (e.g. Martin, 2014). We use results in Table S1 rather than the more general Table 1, because selection coefficients apply to the heterozygous and homozygous effects of alleles in a given background, rather than to the average and dominance effects of substitutions in a population. Note also that the results below apply only with the quadratic fitness function of eq. 1, and not with other fitness functions with higher curvatures that would allow for complex epistasis (i.e. fitness interactions between three or more loci).

To express the results in Table S1 in terms of fitness effects, let us first consider the net effect of evolutionary change – a quantity which corresponds to the fitness effects of whole genotypes. For example, *m*(2**a**, 2**a**) is simply the fitness of one parental genotype, measured in environmental conditions where the alternative parental genotype is optimal:

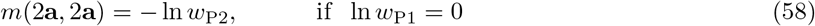

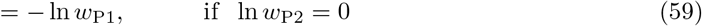

Similarly, *m*(**a** + **d** + **d**) and *m*(**a** – **d**, **a** – **d**) are the fitnesses of the F1 genotype measured in conditions where one or other of the parental genotypes is optimal.

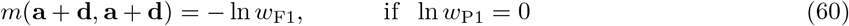

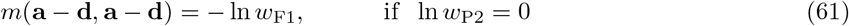

The total amount of evolutionary change depends on the fitness effects of the individual divergent alleles, introgressed one at a time into an optimal background. To see this, let *s_i_* denote the deleterious fitness effect of inserting a single homozygous substitution *i* into an otherwise optimal background. This selection coefficient is defined in the standard way, as *s* = (*w*^1^ – *w*)/*w* where *w′*(*w*) is the fitness of the mutant (wild-type). For small selection coefficients, we also have *s_i_* ≈ – ln(1 – *s_i_*). If the wild-type genotype is phenotypically optimal, it follows that

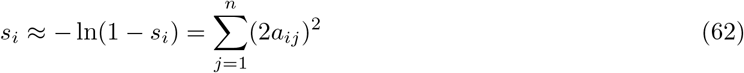

and so, if 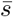 denotes the mean selection coefficient across all *D* substitutions, the total amount of evolutionary change is

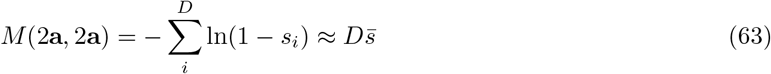

Equivalent results hold for *M*(**a**±**d**, **a**±**d**) for the heterozygous selection coefficients. It follows therefore that the total amount of evolutionary change will be large if the parental lines have fixed many mutations with (potentially) large fitness effects.

We will now show that the difference between the total amount and net effect of change is a measure of fitness epistasis. Let us first note that, with the quadratic model of eq. 1, all epistatic interactions are pairwise (Martin et al., 2007). If we define *s_ik_* as the fitness effect of inserting a given pair of substitutions into an optimal background, then the pairwise epistatic effect is the log fitness of the double mutant, minus the log fitnesses of the two single mutants:

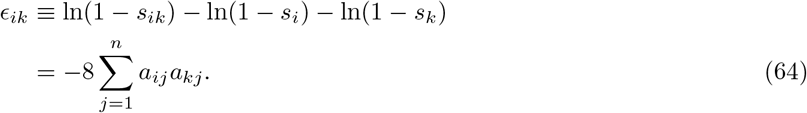

(e.g. Martin et al., 2007). It then follows from eq. 22 that the key quantity for hybrids is

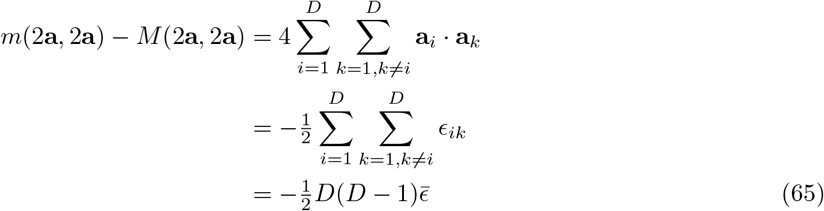

which agrees with results from Chevin et al. (2014). Equation 64 shows that the sign of the fitness epistasis relates to the tendency of mutations to point in the same direction (Martin et al., 2007; Chevin et al., 2014; Fraïsse and Welch, 2019). Deleterious mutations with positive epistasis will tend to be compensatory (pointing in opposite phenotypic directions), and those with negative epistasis will tend to be synergistic (pointing in the same phenotypic direction); epistasis will be maximally negative when all substitutions have identical individual effects, in which case *ϵ* = –2s. Note also that *m*(2**a**, 2**a**) – *M*(2**a**, 2**a**) will vanish when there is no epistasis on average 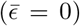, as would be the case if the populations accumulated randomly- orientated mutations (Martin et al., 2007; Simon et al., 2018; Fraïsse and Welch, 2019). Evolutionary differences that show positive epistasis will tend to increase RI among hybrids.

## Appendix 2: Further simulations under stabilizing selection

In this Appendix, we report the results of additional simulations, to explore how the key quantities that determine hybrid fitness (Table 1) behave under stabilizing selection.

### The effects of population genetic parameters under stabilizing selection with the additive model

Let us first consider the effects of varying the population genetic parameters, which have also been explored in several previous studies (Hartl and Taubes, 1996; Poon and Otto, 2000; Welch and Waxman, 2003; Zhang and Hill, 2003; Tenaillon et al., 2007; Lourencço et al., 2011; Chevin et al., 2014; Roze and Blanckaert, 2014; Barton, 2016), but here, we explicitly report the total amount (*M*(**A**, **A**)) and net effect (*m*(**A**, **A**)) of evolutionary change.

To do this, we re-analysed simulation results from Schneemann et al. (2020) each comprised of 500 substitutions accrued under stabilizing selection, with a stationary optimum. Overall, 128 conditions were simulated, using a fully crossed set of parameters. Here, dominance coefficients were drawn from a uniform distribution bounded at 0 and 1, such that mutations were on average phenotypically additive. The parameters varied were (i) the population size (*N* = 1000, or *N* = 10), (ii) the mean selection coefficient of a new mutation in an optimal background (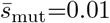 or 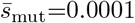), (iii) the genomic mutation rates (*U* ∈ {0.01, 0.001, 0.0001, 0.00001}), (iv) the number of traits under selection (*n* = 2 or *n* = 20), (v) the rate of recombination (either a single chromosome with map length one Morgan, and Haldane’s mapping function, such that the mean crossover fraction was 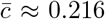; or free recombination among all loci, such that 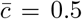), and (vi) the shape of the distribution of mutational effects (either “top down”, where the magnitudes of new mutations were drawn from an exponential distribution, with a random orientation in *n*-dimensional space; or “bottom up”, where the mutational effect on each trait was drawn independently from a normal distribution; Poon and Otto, 2000). Of these six parameters, four had appreciable effects on the results, and these are indicated visually in Figure S1.

The results in Figure S1 show a few clear patterns. First, and unsurprisingly, populations fixed larger changes (larger *M*(**A**, **A**)) when the population size was smaller, and mutations were large (smaller *N*, larger 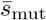). Results for *m*(**A**, **A**) generally support eq. 26, whose value for the four values of *n/N* are shown by the vertical dashed lines (Barton, 2016). The sole exceptions are results with 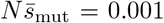 (empty blue points in Fig. S1). In this case, selection was so ineffective that the populations had failed to reach their equilibrium level of maladaptation after *D* = 500 substitutions. In consequence, results fell on the line *m*(**A**, **A**) ≈ *M*(**A**, **A**), implying that the evolutionary changes were wandering erratically in phenotypic space, as under strict neutrality. In all other cases, the action of stabilizing selection was apparent from the fact that *m*(**A**, **A**) ≪ *M*(**A**, **A**).

We note finally that with higher mutation rates the dependencies on *N* and *n* can change (Roze and Blanckaert, 2014). This is due to accumulation of linkage disequilibria, not treated in the current work.

### Dominance effects under stabilizing selection

This section explores stabilizing selection when mutations may be phenotypically dominant or recessive, with a particular focus on the evolution of the dominance effects. In all cases, this will involve modifying the model of mutational dominance reported in the Methods, to enhance the influence of dominance effects.

Let us begin with the simulations reported in Figure 4C&D, which are also reported in greater detail in Figure S2. These simulations used a mutational model of Schneemann et al. (2022). Under this model, as with the standard simulations, the heterozygous effect of a new mutation on a given trait was set to its homozygous effect multiplied by a beta-distributed random number with mean *μ* and variance *ν*. But in this case, both *μ* and *ν* were set to vary with the size of the mutation, such that

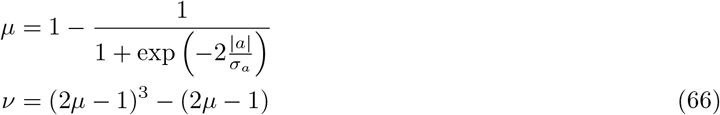

where *σ_a_* is the standard deviation in the additive effects of new mutations. The result is that small-effect mutations were additive on average (with *μ* ≈ 1/2), whereas larger effect mutations became increasingly recessive (Manna et al., 2011; Billiard et al., 2021). Figure S2G (red curve) shows clearly that, with this mutation model, populations evolving under stabilizing selection have a strong tendency to fix phenotypically recessive mutations (eq. 21). Now if P1 had fixed *wholly* recessive mutations (with no phenotypic effect in heterozygous form) then it would follow that *a_ij_* = *d_ij_* for all loci and traits (see Table 5). If we then consider genetically homogeneous parental populations (as in Appendix 1), it would follow trivially that *m*(**A**, **A**) = *m*(**Δ**, **Δ**) = *m*(**A**, **Δ**) and that *M*(**A**, **A**) = *M*(**Δ**, **Δ**) = *M*(**A**, **Δ**). In this way, the tendency for highly recessive mutations to fix, explains the similarities of the red lines shown in Fig. S2C, F and I (which are plotted together in Figure 4D).

**Figure S1:**
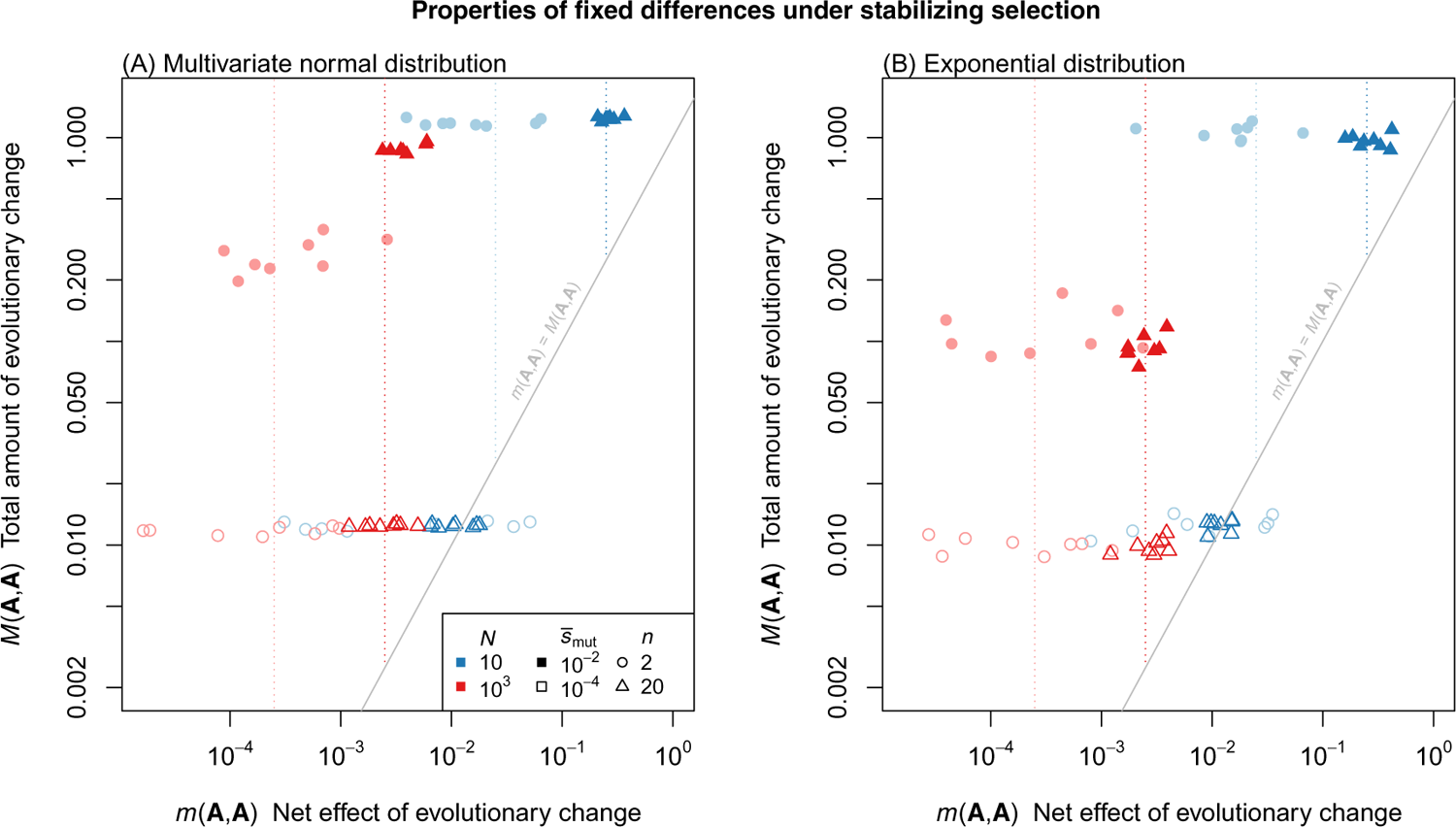
The value for the total amount and net effect of evolutionary change under stabilizing selection depend on model parameters in predictable ways. Simulation results are shown pairs of populations, diverging under stabilizing selection. Simulations used an additive phenotypic model, and were halted after *D* = 500 substitutions have fixed. Each panel contains results from 64 population pairs, using a fully crossed set of population-genetic parameters. Varied were the population size (*N*: red versus blue points), the mean selection coefficient of a new mutation in an optimal background (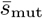: filled versus unfilled points); and the number of phenotypic traits (*n*: circular versus triangular points). Mutation and recombination rates also varied, but neither had a qualitative effect in the parameter regimes simulated, and so are not indicated visually. **(A)** shows results when the mutational effects on each trait were i.i.d. normal. **(B)** shows results when the magnitudes of new mutations were drawn from an exponential distribution, with random orientations in *n*-dimensional space; In both panels, vertical lines show the expected value of *m*(**A**, **A**) at stochastic equilibrium (namely *n*/(8*N*); eq. 26). This equilibrium was not reached, however, when selection was very ineffective (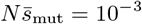: empty blue points), and in this case evolutionary changes wandered erratically in phenotypic space (such that *M*(**A**, **A**) ≈ *m*(**A**, **A**)).

Note, however, that the fixations were not wholly recessive, and so the red lines are similar, but not identical. In particular, a stochastic equilibrium is reached by the red curves in both Figure S2B (eq. 26) and Fig. S2H (where the recessive fixations in P1 imply that the F1 will closely resemble P2: eq. 18). However, from Figure S3E it is clear that the lack of coadaptation between the dominance effects means that their net effect, *m*(**Δ**, **Δ**), still wanders in phenotypic space, and increases steadily with divergence.

**Figure S2:**
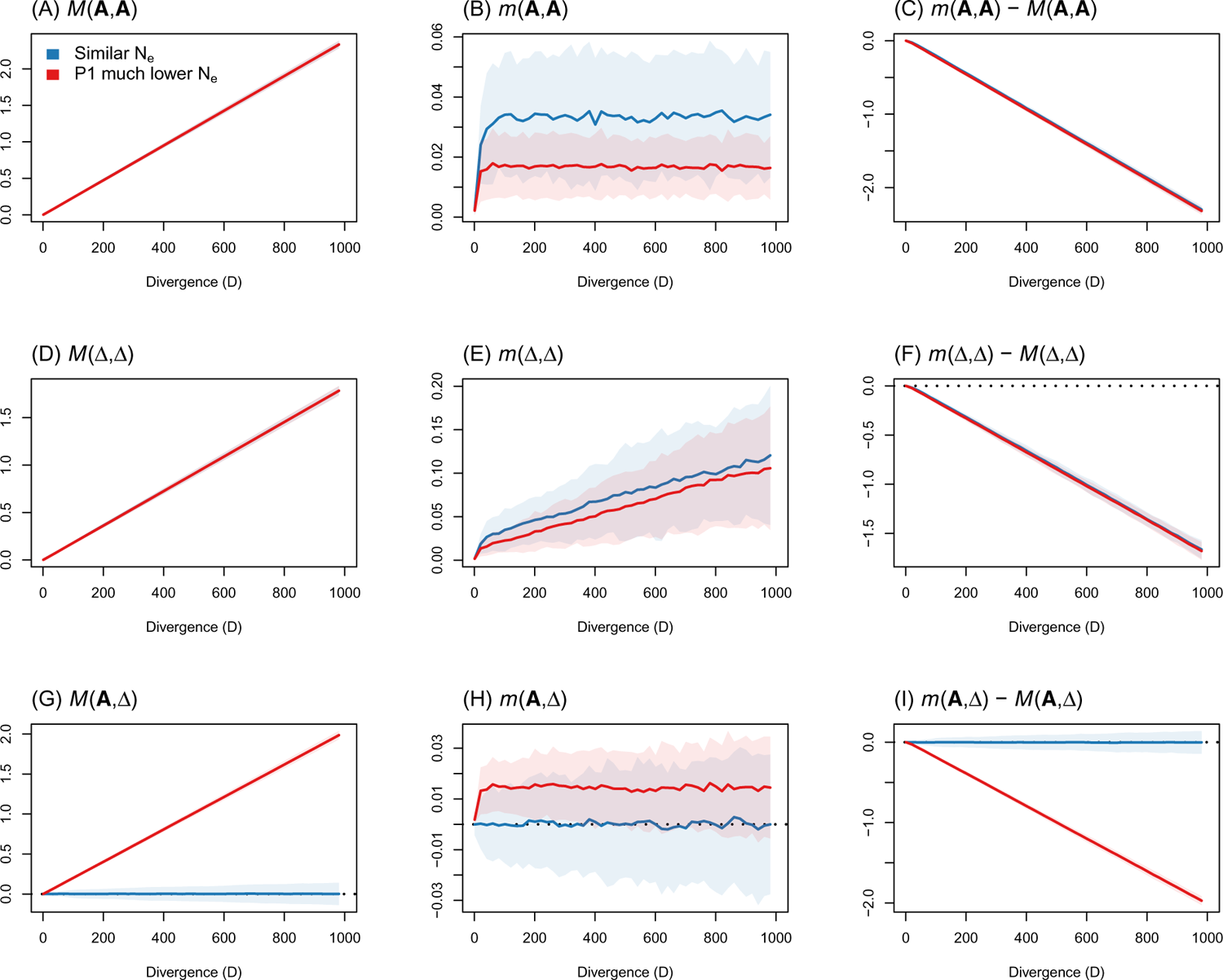
The net effect and total amount of evolutionary change predictably under stabilizing selection, when mutations tend to be phenotypically recessive. The simulations reported correspond to be shown in Figure 4C-D, and the curves in panels C, F and I replicate those in Figure 4C (blue curves), and Figure 4D (red curves). All simulations used the dominance model of Schneemann et al. (2022), in which larger effect mutations were more likely to be phenotypically recessive (eq. 66). All curves show the means across 100 replicate simulations, and shaded areas (often barely visible) show the standard deviation. Other simulation parameters were *N* = 100, *n* = 20 and 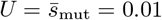.

While the results in Figures 4C-D and S2 assumed that mutations will tend to be phenotypically recessive, it is not clear that this will hold in nature. This is partly because the traits in Fisher’s model need not correspond to real-world quantitative traits (Martin, 2014), and partly because, under the fitness function of eq. 1, mutations can be recessive for fitness, even if they are additive or weakly dominant for the phenotype (e.g. Manna et al., 2011).

As such, we repeated our simulations of stabilizing selection, with no special tendency for mutations to be recessive, but also increasing the variance in the dominance effects. To do this, we simply set *μ* = 1/2 and *ν* = 1/12 so that the heterozygous effect of a new mutant was its homozygous effect, multiplied by a uniformly-distributed random number. As with the main text simulations, we first assumed that each mutation had a unique dominance multiplier on each trait – so that we used *n* uniform random numbers per mutation. However, we also compared this “per-trait dominance” model, to a “per-mutation dominance” model, in which the effects on each trait shared a dominance multiplier – so that we used only a single uniform random number per mutation. The effect of both of these changes to the mutational model was to make it more likely that mutations with extreme levels of dominance would fix, but with no tendency for new mutations to be phenotypically recessive. The results of these simulations are shown Figure S3, with the “per-trait dominance” results as thinner lines, and the “per-mutation dominance” results as thicker lines.

Consider first, results for the interaction terms (Figure S3G-I). Figure S3G shows that a tendency to fix phenotypically recessive mutations (an increasing *M*(**A**, **Δ**)) can occur via a selective sieve without mutational bias, but only for some models of mutation - in this case, only for the “per-mutation” model (thicker red line), in which each mutation has the same level of dominance on all *n* traits. However, the corresponding negative trend in *m*(**A**, **Δ**) – *M*(**A**, **Δ**) (Figure S3I) is now very weak – both compared to its standard deviation between runs (so that the term will be positive for a substantial proportion of runs) – and compared to negative trend in the additive term (Fig. S3C).

Consider finally results for the dominance effects (Figure S3D-F). Remarkably, the trend in Figure S3F is opposite of that shown in Figure S2F, with a weak tend for dominance effects to point in same phenotypic direction. This applies in all cases, including when the sole evolving population tended to fix phenotypically recessive alleles. Note, however, that this tendency is again weak - both compared to its standard deviation and the negative trend in the additive term (Fig. S3C). The upshot is, at least in the models we simulated, dominance terms will be difficult to interpret in the absence of a mutational bias towards phenotypic recessivity.

**Figure S3:**
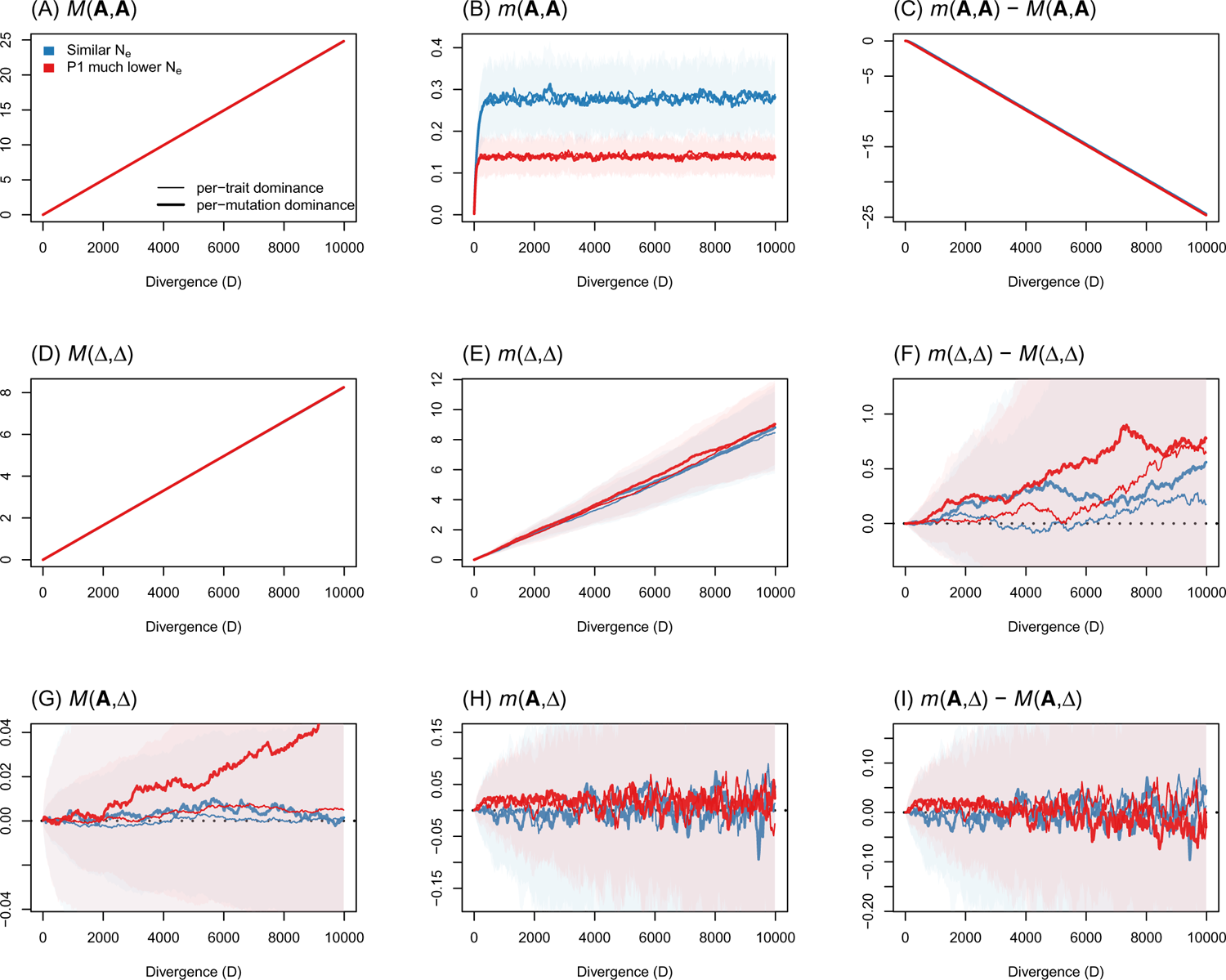
Dominance effects can show weak directionality under stabilizing selection, even without a tendency for mutations to be phenotypically recessive. Simulation results under stabilizing selection, with a stationary optimum. Compared to the main text simulations, the variance in the dominance effects of mutations was increased (by drawing dominance multipliers for each mutation from a uniform distribution with *μ* = 1/2 and *ν* = 1/12), and we also compared our standard model (“per-trait dominance”) to a model in which each mutation was equally dominant or recessive on all n traits (“per-mutation dominance”). Lines and shaded areas represent the mean and one standard deviation across 200 replicate simulations. Other simulation parameters were *N* = 10, *n* = 20 and 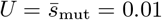

## Acknowledgements

BDS and HS acknowledge support from the Wellcome Trust program in Mathematical Genomics and Medicine (WT220023 and RG92770). We are also very grateful to Matthew Hartfield, Luis-Miguel Chevin, Juan Li, Nick Barton, Roger Butlin, and Anja Westram whose comments greatly improved earlier drafts. This preprint has been peer-reviewed and recommended by Peer Community in Evolutionary Biology at https://doi.org/10.24072/pci.evolbiol.100543.

## Author Contributions

BDS, HS and JJW conceived of the study, BDS and JJW performed the analysis, HS performed the simulations and made the figures, and all authors contributed to writing the manuscript.

## Supporting Information

All scripts and supporting information not given in the appendices can be found at https://github.com/bdesanctis/mode-of-divergence. (release here: https://zenodo.org/badge/latestdoi/533055768) and http://doi.org/10.6084/m9.figshare.21843225.

